# Intranasal pediatric parainfluenza virus-vectored SARS-CoV-2 vaccine candidate is protective in macaques

**DOI:** 10.1101/2022.05.21.492923

**Authors:** Cyril Le Nouën, Christine E. Nelson, Xueqiao Liu, Hong-Su Park, Yumiko Matsuoka, Cindy Luongo, Celia Santos, Lijuan Yang, Richard Herbert, Ashley Castens, Ian N. Moore, Temeri Wilder-Kofie, Rashida Moore, April Walker, Peng Zhang, Paolo Lusso, Reed F. Johnson, Nicole L. Garza, Laura E. Via, Shirin Munir, Daniel Barber, Ursula J. Buchholz

## Abstract

Pediatric SARS-CoV-2 vaccines are needed that elicit immunity directly in the airways, as well as systemically. Building on pediatric parainfluenza virus vaccines in clinical development, we generated a live-attenuated parainfluenza virus-vectored vaccine candidate expressing SARS-CoV-2 prefusion-stabilized spike (S) protein (B/HPIV3/S-6P) and evaluated its immunogenicity and protective efficacy in rhesus macaques. A single intranasal/intratracheal dose of B/HPIV3/S-6P induced strong S-specific airway mucosal IgA and IgG responses. High levels of S-specific antibodies were also induced in serum, which efficiently neutralized SARS-CoV-2 variants of concern. Furthermore, B/HPIV3/S-6P induced robust systemic and pulmonary S-specific CD4^+^ and CD8^+^ T-cell responses, including tissue-resident memory cells in lungs. Following challenge, SARS-CoV-2 replication was undetectable in airways and lung tissues of immunized macaques. B/HPIV3/S-6P will be evaluated clinically as pediatric intranasal SARS-CoV-2/parainfluenza virus type 3 vaccine.

**One-Sentence Summary:** Intranasal parainfluenza virus-vectored COVID-19 vaccine induces anti-S antibodies, T-cell memory and protection in macaques.

## INTRODUCTION

Severe acute respiratory syndrome coronavirus 2 (SARS-CoV-2) infects and causes disease in all age groups. Although COVID-19 is generally milder in young children than in adults, thousands of children have been hospitalized due to COVID-19 in the US alone, including about one-third without preexisting medical conditions (Funk et al., 2022; Rankin et al., 2021). More than 800 children 0 to 11 years of age have died from COVID-19 in the US (https://covid.cdc.gov/covid-data-tracker/#demographics, accessed on April 22, 2022), and during the fall/winter COVID-19 wave of 2021/2022, children accounted for over 25% of US cases (Gerber and Offit, 2021). In rare cases (∼0.03% of infected children), COVID-19 can cause a multisystem inflammatory syndrome in children [MIS-C; (Riphagen et al., 2020; Verdoni et al., 2020)], arising within about 4 weeks after SARS-CoV-2 exposure. While mRNA-based vaccines are available for children 5 years of age and older, to date, no vaccine has been authorized or recommended for children under 5 years of age. Furthermore, interim results from an ongoing Phase 1/2/3 clinical study suggested that a third 3-µg dose of the BNT162b2 mRNA vaccine may be needed to elicit robust immune responses in children >2 to <5 years of age; accordingly, a 3-dose regimen is being evaluated in children >6 months to <5 years of age (NCT04816643). One limitation of the current mRNA and other injectable SARS-CoV-2 vaccines is that they do not directly stimulate immunity in the respiratory tract, the major site of SARS-CoV-2 entry, replication, disease, and egress (DiPiazza et al., 2021). Therefore, it is important to evaluate additional vaccine approaches that are suitable for pediatric use and stimulate both mucosal and systemic immunity. Ideally, a vaccine should be effective at a single dose, and could be administered topically to the respiratory tract to induce robust systemic and respiratory mucosal immunity that restricts SARS-CoV-2 infection and shedding.

Here, we describe the pre-clinical evaluation of the safety, immunogenicity, and protective efficacy of a live intranasal SARS-CoV-2 vaccine candidate, B/HPIV3/S-6P, in rhesus macaques. B/HPIV3/S-6P consists of a live-attenuated chimeric bovine/human parainfluenza virus type-3 (B/HPIV3) that was modified to express the SARS-CoV-2 S-protein trimer stabilized in the prefusion form. The B/HPIV3 vector originally was developed as a pediatric vaccine candidate against human PIV3 (HPIV3), a single-stranded negative-sense RNA virus which is an important cause of respiratory illness, especially in infants and young children under 5 years of age (DeGroote et al., 2020; Howard et al., 2021). Previously, B/HPIV3 was well-tolerated in a Phase 1 study in infants and young children (Karron et al., 2012). B/HPIV3 also has been used to express the fusion (F) glycoprotein of another human respiratory pathogen, human respiratory syncytial virus (RSV), providing a bivalent HPIV3/RSV vaccine candidate which was well-tolerated in children >2 months of age [(Bernstein et al., 2012), Clinicaltrials.gov NCT00686075]. We recently reported that B/HPIV3 expressing a stabilized prefusion form of S efficiently protected hamsters against infection with a vaccine antigen-matched SARS-CoV-2 isolate, preventing weight loss, lung inflammation and efficiently reducing SARS-CoV-2 replication in the upper and lower respiratory tract (Liu et al., 2021). In the present study, we evaluated the safety and efficacy of a single intranasal/intratracheal (IN/IT) dose of B/HPIV3/S-6P in rhesus macaques (RM). Immunogenicity evaluations included S-specific mucosal and systemic antibody and T-cell responses as well as neutralizing antibody responses to the vaccine-matched SARS-CoV-2 strain and four major variants of concern. In addition, we assessed the protective efficacy of B/HPIV3/S-6P against SARS-CoV-2 challenge prior to advancing this candidate to a Phase 1 clinical study.

## RESULTS

### B/HPIV3/S-6P replicates in the upper and lower airways of rhesus macaques

We used B/HPIV3 to express a prefusion-stabilized version of the SARS-CoV-2 S protein. B/HPIV3 is a cDNA-derived version of bovine PIV3 (BPIV3) strain Kansas in which the BPIV3 hemagglutinin-neuraminidase (HN) and fusion (F) glycoproteins (the two PIV3 neutralization antigens) have been replaced by those of the human PIV3 strain JS (Karron et al., 2012; Schmidt et al., 2000) (Figure 1A). The bovine PIV3 backbone provides stable attenuation due to host-range restriction in humans (Bernstein et al., 2012; Karron et al., 2012). B/HPIV3/S-6P expresses a full-length prefusion-stabilized version (S-6P) of the SARS-CoV-2 S protein (1,273 aa) from a supernumerary gene, inserted between the N and P genes, an insertion site previously shown to be optimal for efficient expression and genetic stability (Liang et al., 2014) (Figure 1A). The S-6P version of the S-protein contains 6 proline substitutions (Hsieh et al., 2020) that stabilize S in its trimeric prefusion form and increase expression and immunogenicity. The S1/S2 polybasic furin cleavage motif “RRAR” was ablated by amino acid substitutions (RRAR-to-GSAS) (Wrapp et al., 2020) (Figure 1A), rendering S-6P non-functional for virus entry, which eliminates the possibility of S altering the tissue tropism of the B/HPIV3 vector.

**Figure 1.**
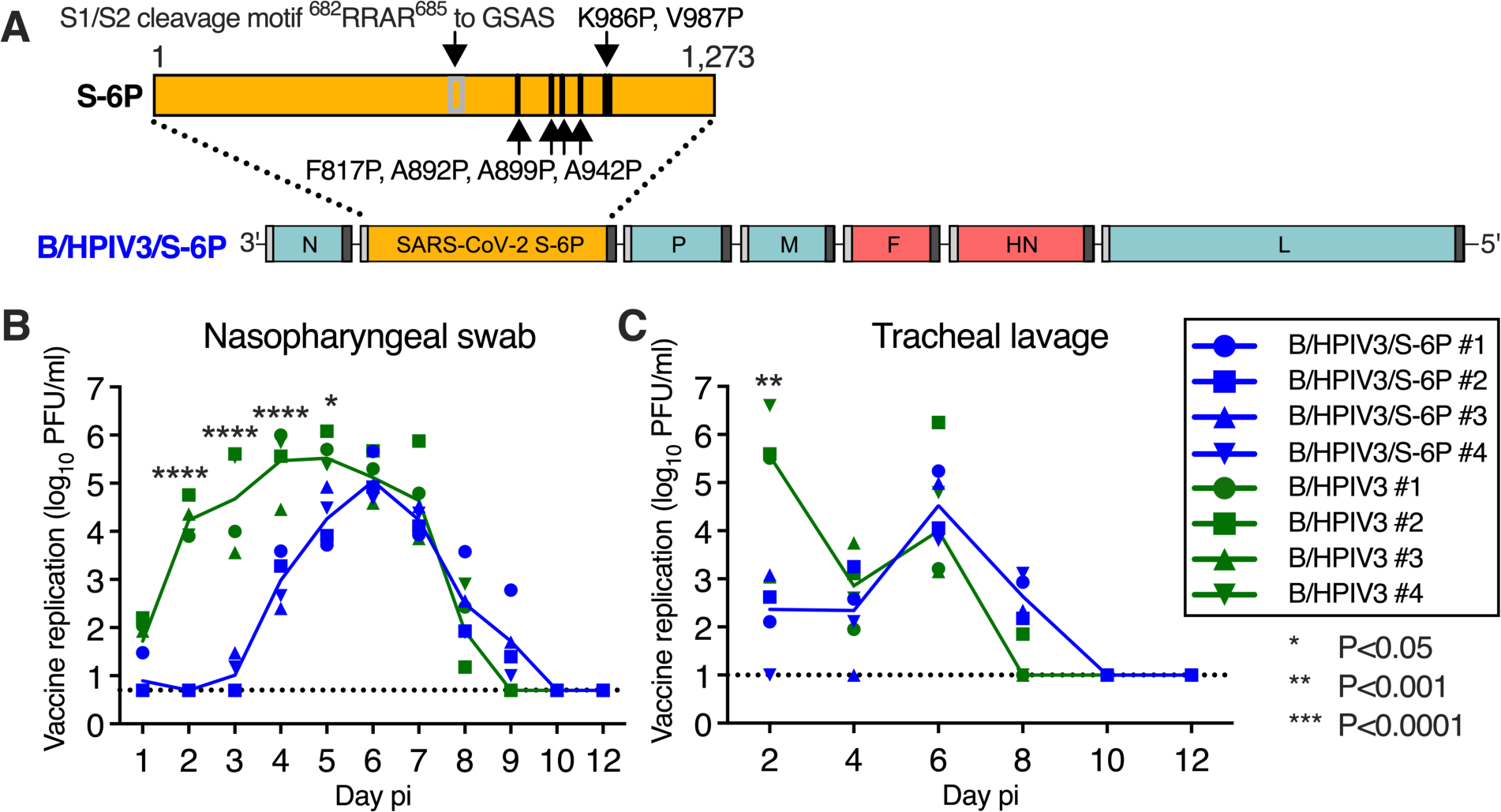
Genome organization of B/HPIV3/S-6P; vaccine replication following intranasal/intratracheal immunization of rhesus macaques. (**A**) Diagram of the B/HPIV3/S-6P genome, with BPIV3 (N, P, M and L; blue) and HPIV3 genes (F and HN; red). The full-length SARS-CoV-2 S ORF (codons 1-1,273) was codon-optimized and inserted as additional gene (orange) between the N and P ORFs. The S sequence includes 6 stabilizing proline substitutions (S-6P) and RRAR-to-GSAS substitutions to ablate the S1/S2 cleavage site. Each gene begins and ends with PIV3 gene-start and gene-end transcription signals (light and dark bars). (**B** and **C**) Replication of B/HPIV3/S-6P and B/HPIV3 in upper (**B**) and lower (**C**) airways of rhesus macaques (RM). Two groups of 4 RMs were immunized intranasally and intratracheally with 6.3 log_10_ PFU of B/HPIV3/S-6P (blue) or B/HPIV3 (green). Nasopharyngeal swabs and tracheal lavages were performed daily and every other day, respectively, on days 0 to 12 post-immunization (pi). Vaccine virus titers were determined by immunoplaque assay (Materials and Methods); expressed as log_10_ PFU/ml [Limit of detection: 0.7 log_10_ PFU/mL for nasopharyngeal swabs; 0.7 log_10_ PFU/mL for tracheal lavages (dotted line)]. Each RM is indicated by a symbol; lines represent medians (*P<0.05, **P<0.01, ****P<0.0001; two-way ANOVA, Sidak multiple comparison test).

To evaluate vaccine replication and immunogenicity, we immunized 2 groups of RMs (n=4 per group) with a single dose of 6.3 log_10_ plaque-forming units (PFU) of B/HPIV3/S-6P or the B/HPIV3 vector control, respectively, administered by the combined intranasal and intratracheal route (IN/IT) (Figure S1). Nasopharyngeal swabs (NS) and tracheal lavages (TL) were performed daily and every other day, respectively, from day 0 to 12 post-immunization (pi) to evaluate vaccine replication in the upper and lower airways (UA and LA, respectively; Figure 1B-C, Figure S1). Replication of B/HPIV3/S-6P and the B/HPIV3 control was detectable through days 8 or 9 pi in the UA and LA. In the UA, peak replication of B/HPIV3/S-6P and B/HPIV3 control was detected between study days 4 and 6 (medians independent of study day: 4.9 log_10_ PFU/mL vs 5.9 log_10_ PFU/mL, respectively; P=0.1429 by two-tailed Mann-Whitney test); replication of B/HPIV3/S-6P was delayed by 1-2 days compared to that of the empty vector (P<0.0001 on days 2 to 4) (Figure 1B). In the LA, B/HPIV3/S-6P replicated with similar kinetics as B/HPIV3, reaching median peak titers of 4.5 log_10_ PFU/mL and 4.0 log_10_ PFU/mL, respectively, on day 6 pi (Figure 1C).

To evaluate the stability of S expression during vector replication, NS and TL specimens positive for B/HPIV3/S-6P were evaluated by a dual-staining immunoplaque assay, which detects the expression of S and vector proteins. On average, 89% of the B/HPIV3/S-6P plaques recovered between days 5 and 7 from NS were positive for S expression (Figure S2), suggesting stable S-6P expression in the UA. In TL specimens collected on day 6 pi, S expression was stable in 3 of 4 RM, with on average 88% of the plaques positive for S expression. In TL samples from one B/HPIV3/S-6P-immunized RM (B/HPIV3/S-6P #4), plaques were negative for S expression on day 6 pi. Sanger sequencing of the S gene revealed 13 cytidine-to-thymidine mutations in a 430-nucleotide region, suggestive of deaminase activity in the LA of this animal. Eleven were missense mutations resulting in amino acid substitutions, including 7 proline substitutions which might affect S protein folding.

No changes in body weight, rectal temperature, respiration, oxygen saturation or pulse were detected following immunization of RMs with B/HPIV3 or B/HPIV3/S-6P (Figure S3). Thus, B/HPIV3/S-6P replicates efficiently in the UA and LA of RMs, causes no apparent symptoms, and is cleared in approximately ten days.

### B/HPIV3/S-6P induces anti-SARS-CoV-2 S mucosal antibodies in the upper and lower airways

To assess the kinetics of airway mucosal antibody responses to the SARS-CoV-2 S protein in the UA and LA, we collected nasal washes (NW) 3 days before immunization and on days 14, 21, and 28 after immunization, and bronchoalveolar lavage (BAL) fluid on days 9, 21, and 28 pi (Figure S1). We evaluated IgA and IgG binding antibodies using a soluble prefusion stabilized S-2P version of the vaccine-matched S protein (Wrapp et al., 2020) or its receptor binding domain (RBD) (Wrapp et al., 2020) in a highly-sensitive dissociation-enhanced lanthanide fluorescence (DELFIA) immunoassay (Figures 2, A and B).

**Figure 2.**
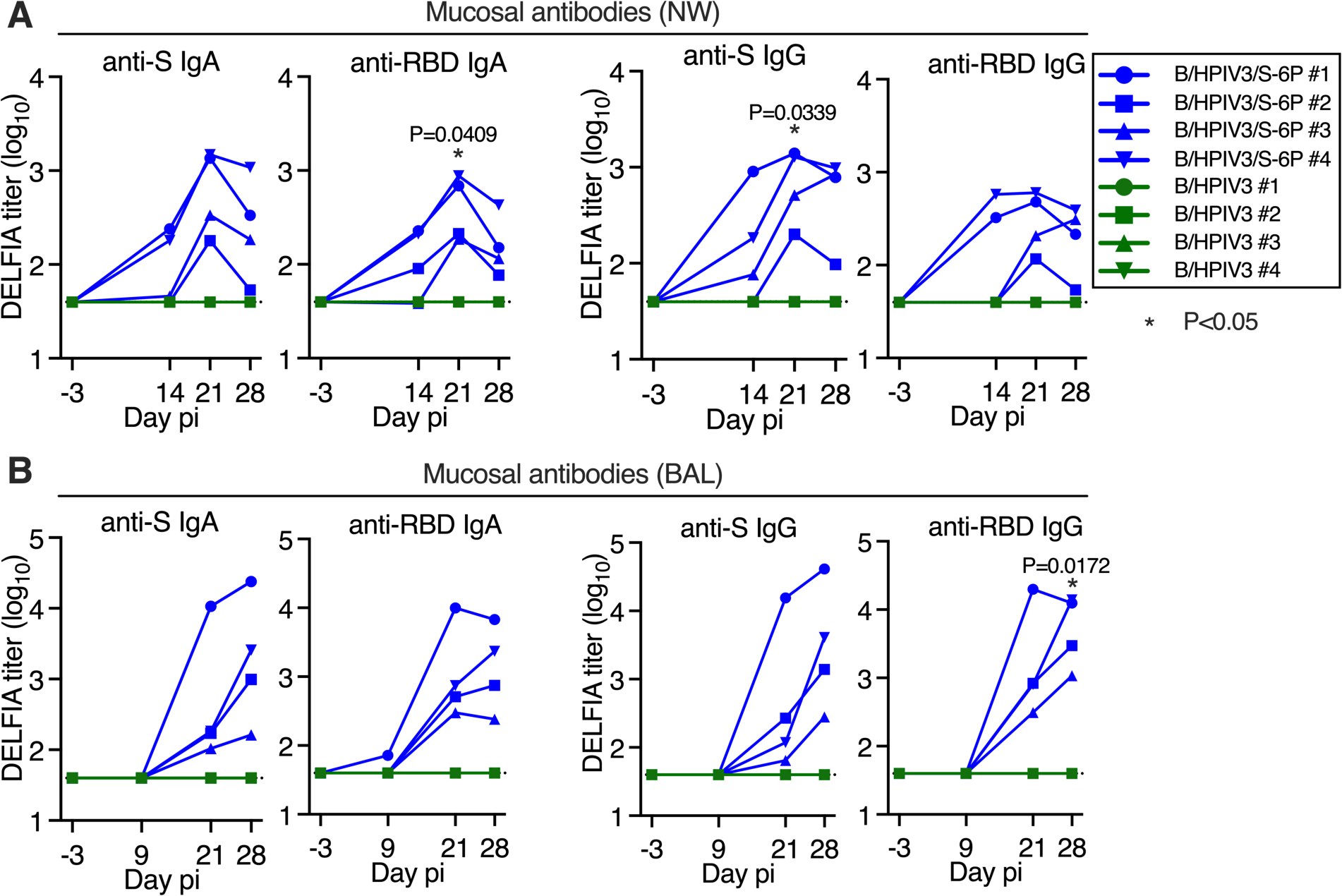
Intranasal/intratracheal immunization with B/HPIV3/S-6P induces mucosal antibody responses to SARS-CoV-2 S in the upper and lower airways. Rhesus macaques (n=4 per group) were immunized with B/HPIV3/S-6P or B/HPIV3 (control) by the intranasal/intratracheal route (Figure S1). To determine the mucosal antibody response in the upper airways, nasal washes (NW) were performed before immunization and on days 14, 21, and 28. To analyze the antibody response in the lower airways, bronchoalveolar lavages (BAL) were collected before immunization and on days 9, 21, and 28 pi. (**A** and **B**) S- and receptor binding domain (RBD)-specific mucosal IgA and IgG titers on indicated days post-immunization (pi) in the upper (A) and lower (B) airways, determined by time-resolved dissociation-enhance lanthanide fluorescence (DELFIA-TRF) immunoassay. Endpoint titers are expressed in log_10_ for mucosal IgA and IgG to a secreted prefusion-stabilized form (aa 1-1,208; S-2P) of the S protein (left panels) or to a fragment of the S protein (aa 328-531) containing SARS-CoV-2 RBD (right panels). The limit of detection is 1.6 log_10_ (dotted line). B/HPIV3/S-6P-immunized RMs are shown in blue, while B/HPIV3-immunized RMs are in green, with each RM represented by a symbol. *P<0.05 (two-way ANOVA, Sidak multiple comparison test).

In B/HPIV3/S-6P-immunized animals, we detected mucosal anti-S (2/4 animals) and anti-RBD IgA (3/4 animals) in the UA as early as 14 days pi (Figure 2A). By day 21 pi, all 4 B/HPIV3/S-6P immunized RMs exhibited anti-S and anti-RBD IgA (geometric mean titers (GMT) between 2.3 and 3.2 log_10_, P=0.0409 for anti-RBD IgA on day 21 pi). B/HPIV3/S-6P also induced mucosal anti-S and anti-RBD IgG responses in the UA on day 14 pi in 3/4 and 2/4 RMs, respectively, with responses observed in all animals by day 21 (anti-S titers between 2.1 and 3.1 log_10_, P=0.0339).

B/HPIV3/S-6P also induced mucosal anti-S and anti-RBD IgA and IgG in the LA (Figure 2B). On day 21 pi, anti-S and anti-RBD IgA titers between 2.0 and 4.0 log_10_ were detectable in the LA of all 4 B/HPIV3/S-6P-immunized RMs. Anti-S IgA titers in the LA continued to rise in all RMs until day 28 after immunization. On day 21 pi, all B/HPIV3/S-6P-immunized RMs also had anti-S and anti-RBD IgG detectable (GMT between 1.8 and 4.2 log_10_). Anti-S IgG titers continued to rise between days 21 and 28 pi. Similarly, anti-RBD IgG titers continued to rise until day 28 pi in 3 RMs, but modestly declined in 1 RM. As expected, none of the RMs immunized with the empty B/HPIV3 vector had detectable anti-S or anti-RBD IgA or IgG antibodies.

### B/HPIV3/S-6P induces serum antibodies against SARS-CoV-2 S that neutralize SARS-CoV-2 WA1/2020 and variants of concern (VoCs)

We next assessed the kinetics and breadth of the serum antibody response to B/HPIV3/S-6P (Figure 3). We detected robust serum IgM, IgA and IgG binding antibody responses to the S protein and RBD by ELISAs in 4/4 B/HPIV3/S-6P-immunized RMs as early as 14 days pi (Figure 3A). Serum anti-S and anti-RBD IgM titers peaked on day 21 pi in all 4 RMs (titers between 4.1 and 5.3 log_10_, P<0.05), and declined towards day 28 pi. Serum anti-S IgA titers peaked on day 21 pi in 2 RMs and remained steady, while they continued to rise until day 28 pi in the other 2 RMs (peak titers between 4.3 and 4.9 log_10_, P<0.01). Serum anti-RBD IgA titers peaked on day 21 in all 4 RMs (titers between 4.8 and 5.3 log_10_, P<0.01) and modestly declined by day 28 pi. High levels of serum anti-S and anti-RBD IgG were also measured in all B/HPIV3/S-6P-immunized RMs on day 14 pi, continuing to rise in all RMs until day 28 pi (GMTs between 5.8 to 6.4 log_10_ on day 28 pi, P<0.0001). These levels of anti-S and anti-RBD IgG antibodies were 16-fold and 180-fold higher than the mean anti-S and anti-RBD IgG titers, respectively, detected in the plasma obtained from 23 SARS-CoV-2-convalescent humans. As expected, 0/4 RMs immunized with empty B/HPIV3 control had serum anti-S or anti-RBD IgM, IgA or IgG antibodies detectable at any time.

**Figure 3.**
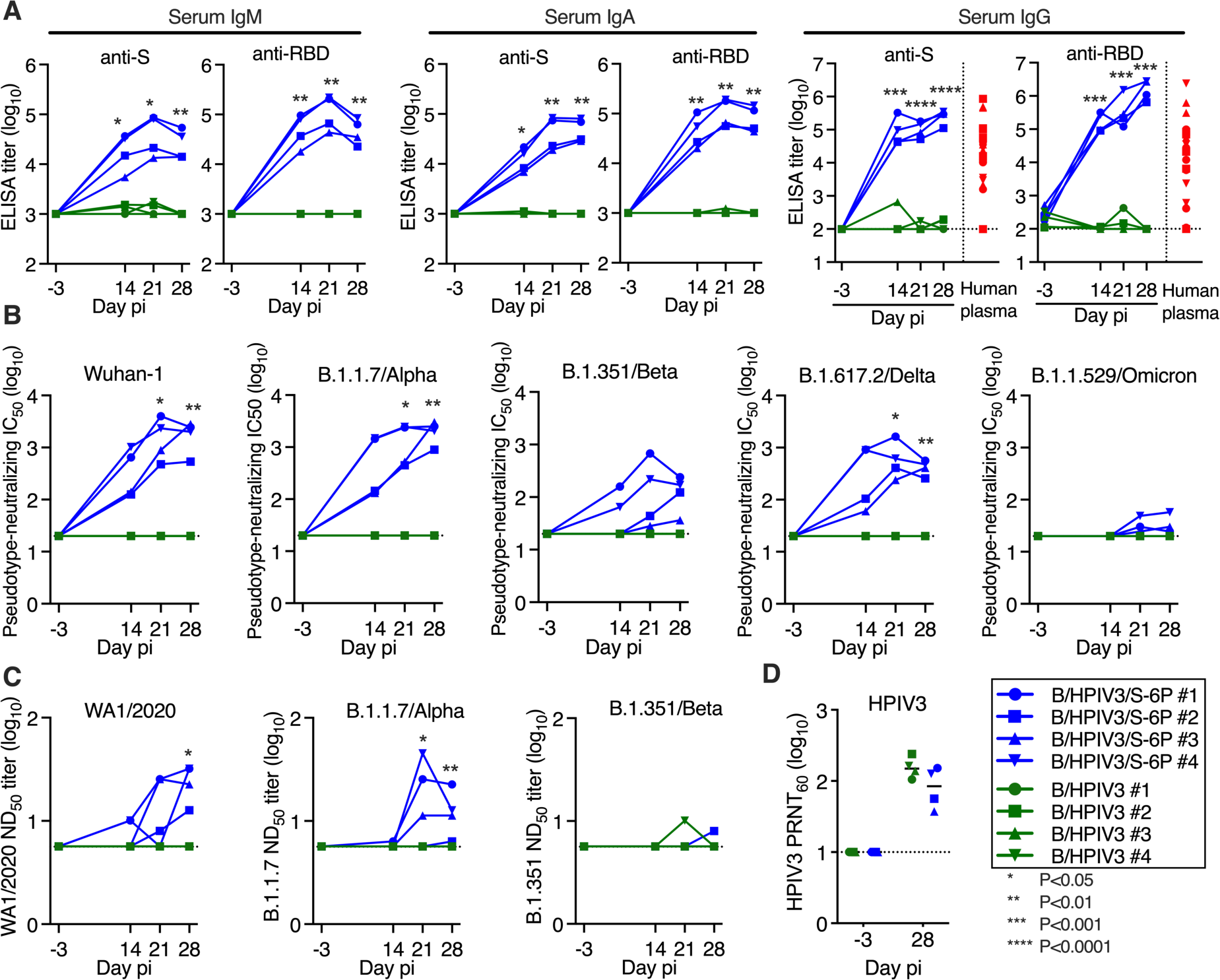
B/HPIV3/S-6P induces serum binding antibody responses to SARS-CoV-2 S and neutralizing antibody responses to VoCs in RMs. Sera were collected from RMs before immunization and on days 14, 21, and 28 pi. (**A**) Endpoint ELISA titers of serum IgM, IgA and IgG to S-2P (left panels) or to the receptor binding domain (RBD, right panels), expressed in log_10_. Twenty-three plasma samples from COVID-19 convalescent individuals were evaluated in parallel for serum IgG to S-2P or the RBD (red symbols). The limits of detection are 3 log_10_ for IgM and IgA and 2.0 log_10_ for IgG. (**B**) Serum neutralizing titers to pseudoviruses bearing spike proteins from SARS-CoV-2 Wuhan-1 (matching S-6P) (Wrapp et al., 2020), B.1.1.7/Alpha, B.1.351/Beta, B.1.617.2/Delta or B.1.1.529/Omicron. The 50% inhibitory concentration (IC_50_) titers of sera were determined. The detection limit is 1.3 log_10_. (**C**) The 50% SARS-CoV-2 serum neutralizing titers (ND_50_) were determined on Vero E6 cells against vaccine-matched WA1/2020, or viruses from lineages B.1.1.7/Alpha or B.1.351/Beta. The limit of detection is 0.75 log_10_. (**D**) Serum HPIV3 neutralizing antibody titers, determined by 60% plaque reduction neutralization test (PRNT_60_); pi, post-immunization.The detection limit is 1 log_10_. Each RM is represented by a symbol. *P<0.05, **P<0.01, ***P<0.001, ****P<0.0001, 2-way-ANOVA, Sidak multiple comparison test.

We also evaluated the kinetics and breadth of the serum neutralizing antibody response to vaccine-matched SARS-CoV-2 and to 4 VoCs (B.1.1.7/Alpha, B.1.351/Beta, B.1.617.2/Delta, and B.1.1.529/Omicron BA.1 sublineage) using a lentivirus-based pseudotype neutralization assay (Corbett et al., 2020a) (Figure 3B). The sera efficiently and similarly neutralized lentivirus pseudotyped with vaccine-matched Wuhan-1 S protein (IC_50_ on day 28 between 2.7 and 3.5 log_10_) or with S from the Alpha lineage (IC_50_ between 3.0 and 3.5 log_10_). The sera also neutralized the Beta S-pseudotyped lentivirus, although the titer was reduced compared to the vaccine match (IC_50_ between 1.6 and 2.4 log_10_). Day 14 sera from all 4 RMs efficiently neutralized the Delta S-pseudotyped lentivirus; titers further increased, but, on day 28, were about 5-fold reduced compared to the vaccine match (IC_50_ between 2.4 and 2.8 log_10_). A low neutralizing activity against Omicron BA.1 was detected in day 28 sera from 3 of 4 RMs (IC_50_ between 1.4 and 1.8 log_10_) that was 59-fold reduced compared to the vaccine match.

The serum neutralizing antibody titers were also assessed by a live-virus SARS-CoV-2 neutralization assay using the vaccine-matched WA1/2020 isolate or an isolate of the Alpha or Beta lineages (Figure 3C). Results were overall comparable with those of the pseudotyped lentivirus neutralization assays, although, as expected, the sensitivity and the dynamic range of the live virus neutralization assays were lower than those of the pseudotype neutralization assays. As expected, neutralizing antibodies against the various SARS-CoV-2 lineages were undetectable in sera from B/HPIV3-control immunized RMs by pseudotype or SARS-CoV-2 neutralization assay; additionally, all 8 RMs developed neutralizing serum antibodies against the HPIV3 vector (PRNT_60_ titers between 1.6 and 2.4 log_10_, Figure 3D).

### B/HPIV3/S-6P immunization induces high frequencies of SARS-CoV-2 S-specific CD4^+^ and CD8^+^ T-cells in the blood and the airways

SARS-CoV-2 S-specific CD4^+^ and CD8^+^ T-cell responses were evaluated using peripheral blood mononuclear cells (PBMCs) and cells recovered from the LA by BAL (see Figure S4 for the gating strategy) at the indicated time points following immunization with B/HPIV3/S-6P (Figs. 4 and 5, figs. S5 and S6) and challenge with SARS-CoV-2 (Figure 4, Figure S7). SARS-CoV-2 S and N-specific CD4^+^ and CD8^+^ T-cells were identified as IFNγ^+^/TNFα^+^ double-positive cells after stimulation with pools of overlapping 15-mer peptides covering the entire length of these proteins. S-specific CD4^+^ T-cells were present in the blood of all B/HPIV3/S-6P-immunized RMs by day 9 pi (Figure 4A, left panels; kinetics are shown in Figure 4B); frequencies peaked on day 9 (2 RMs) or day 14 pi (2 RMs; average peak % of S-specific CD4^+^ T-cells irrespective of the peak day of 0.6%), and then steadily declined until day 28 pi. S-specific CD8^+^ T-cells were also detectable in the blood of B/HPIV3/S-6P-immunized RMs on days 9 pi (Figure 4A, right panels, and Figure 4C), and their frequencies peaked on day 14 pi in 3 of 4 RMs (Figure 4C; average peak % of S-specific CD8^+^ T-cells irrespective of the peak day of 1.1%).

**Figure 4.**
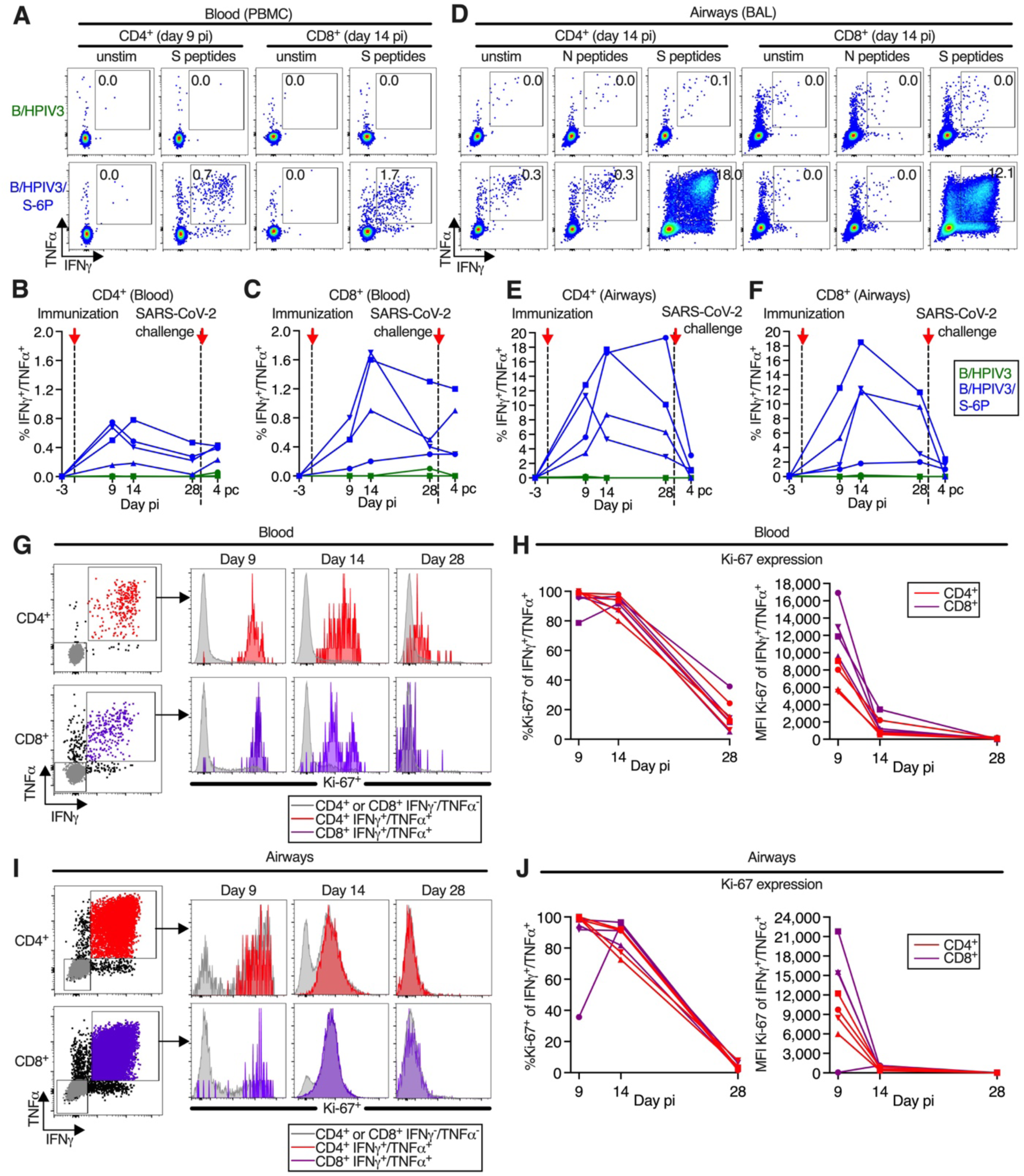
Intranasal/intratracheal immunization with B/HPIV3/S-6P induces S-specific CD4^+^ and CD8^+^ T-cell responses in blood and lower airways. (**A** to **F**) Frequencies of S-specific CD4^+^ and CD8^+^ T cells from blood (A to C) or BAL (D to F). Mononuclear cells collected on indicated days post-immunization (pi) were stimulated with overlapping SARS-CoV-2 S or (BAL only) N peptides or left unstimulated, and processed for flow cytometry. Phenotypic analyses were performed on non-naïve non-regulatory (CD95^+^/Foxp3^-^) CD4^+^ or CD8^+^ T-cells (see Figure S4 for gating); frequencies are relative to that population. (A and D) Dot plots showing IFNγ and TNFα expression by CD4^+^ or CD8^+^ T-cells from blood (A) or BAL (D) of representative B/HPIV3 (top) or B/HPIV3/S-6P-immunized (bottom) RMs. (B, C, E, F) Background-corrected frequencies of S-specific IFNγ^+^/TNFα^+^ CD4^+^ (C, E) or CD8^+^ (D, F) T-cells from blood (C, D) or BAL (E, F) on indicated days. (**G** to **J**) Expression of proliferation marker Ki-67 by IFNγ^+^/TNFα^+^ CD4^+^ (red) or CD8^+^ (purple) T-cells from blood (G and H) or from BAL (I and J) of B/HPIV3/S-6P-immunized RM (n=4, each represented by different symbols). IFNγ^-^/TNFα^-^ cells in grey. (G and I) Gating and histograms showing Ki-67 expression. (H and J) Frequencies of Ki-67^+^ T-cells and median fluorescence intensity (MFI) in IFNγ^+^/TNFα^+^ T-cells from blood (H) or BAL (J). BAL, bronchoalveolar lavage.

**Figure 5.**
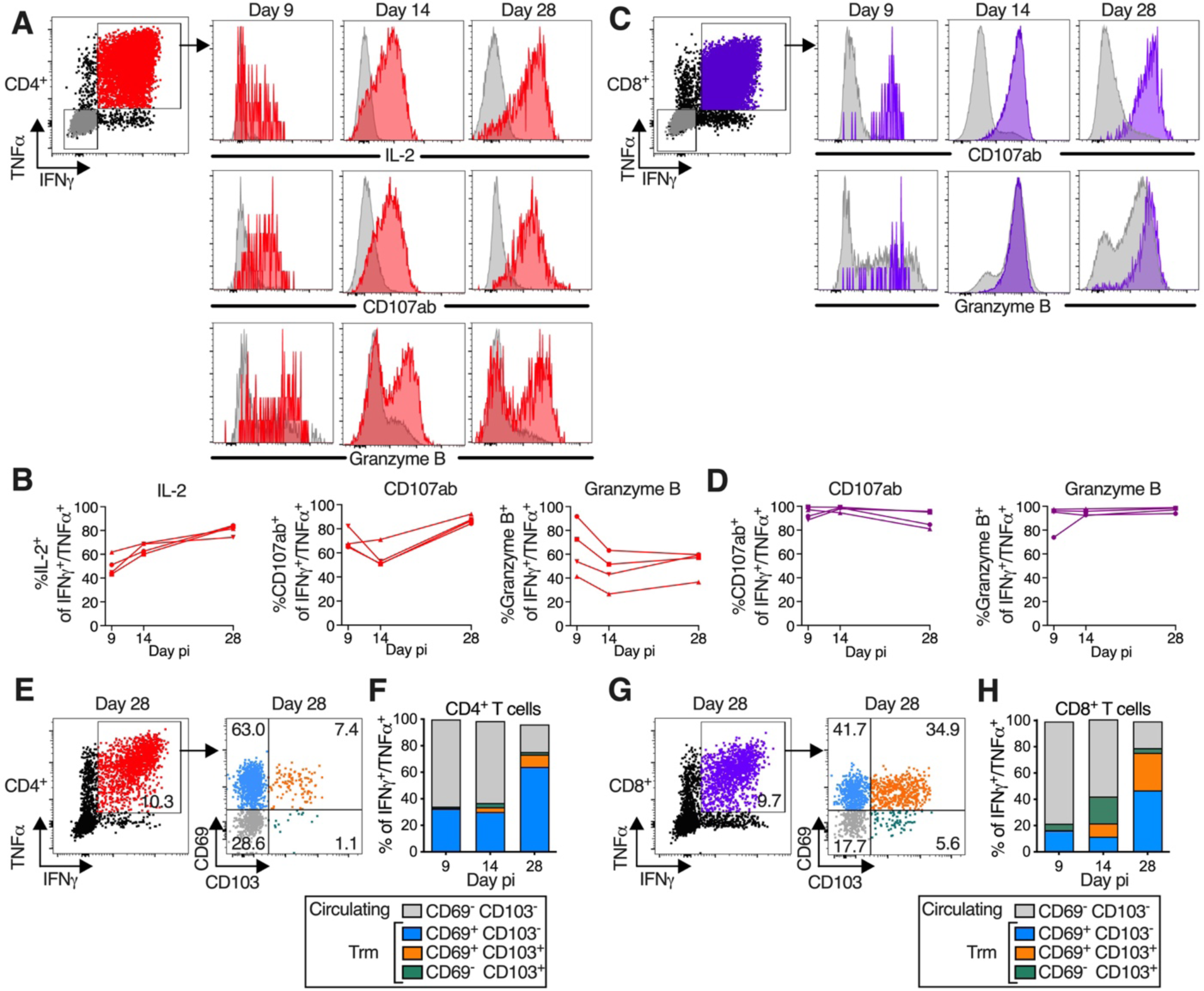
B/HPIV3/S-6P-elicited S-specific CD4^+^ and CD8^+^ T-cells in lower airways (LA) transition to tissue-resident memory phenotype. T-cells obtained by bronchoalveolar lavage (BAL), stimulated with overlapping S peptides prior to flow cytometry analysis. (**A** to **D**) S-specific T-cells in LA are functional. (A and C) Representative dot plots showing gating on S-specific IFNγ^+^/TNFα^+^ CD4^+^ (red), CD8^+^ (purple), and IFNγ^-^/TNFα^-^ (grey) T-cells (for complete gating, see Figure S4); histograms showing expression of IL-2 (CD4^+^ T-cells only), CD107ab and granzyme B by IFNγ^+^/TNFα^+^ T-cells collected on indicated days p.i.. (B and D) Frequencies of IL-2^+^, CD107ab^+^ and granzyme B^+^ of IFNγ^+^/TNFα^+^ S-specific CD4^+^ (B) or CD8^+^ (D) T-cells from 4 BHPIV3/S-6P-immunized RMs on indicated days. (**E** to **H**) Transition to memory phenotype. (E and G) Representative dot plots showing gating on S-specific IFNγ^+^/TNFα^+^ CD95^+^/Foxp3^-^ T-cells (left panels). CD69 and CD103 were used to differentiate circulating (CD69^-^/CD103^-^, grey) and tissue-resident memory [Trm; CD69^+^/CD103^-^ (blue), CD69^+^/CD103^+^ (orange), CD69^-^/CD103^+^ (green)] S-specific IFNγ^+^/TNFα^+^ T-cells from LA (right panels, % indicated). (F, H) The median % of circulating and each of the 3 Trm S-specific IFNγ^+^/TNFα^+^ CD4^+^ (F) or CD8^+^ (H) T-cell subsets present on indicated days in BAL of 4 B/HPIV3/S-6P-immunized RMs are stacked.

In the LA of B/HPIV3/S-6P-immunized animals, S-specific CD4^+^ and CD8^+^ T-cells were abundant by day 9 pi or 14 (Figure 4, D to F). Remarkably, the average peak percentage of S-specific CD4^+^ T-cells recovered from BAL irrespective of day pi reached 14.3% (Figure 4E). In 3 of 4 animals, their frequency declined between day 14 and 28 pi. S-specific CD8^+^ T-cells in BAL also peaked on day 14 pi in 3 of 4 RMs (Figure 4F; average peak % of S-specific IFNγ^+^/TNFα^+^ CD8^+^ T-cells irrespective of the peak day of 11.1%). No S-specific CD4^+^ or CD8^+^ T-cells were detected in the blood or BAL of RMs immunized with B/HPIV3 (Figure 4, A to F). Lastly, stimulation with SARS-CoV-2 N peptides of CD4^+^ or CD8^+^ T-cells isolated from BAL, which was included as negative control, did not reveal IFNγ^+^/TNFα^+^ positive cells above the background present in unstimulated cells (Figure 4D).

On day 9 pi, close to 100% of the S-specific CD4^+^ and CD8^+^ T-cells in the blood (Figure 4, G and H) and BAL (Figure 4, I and J) of the B/HPIV3/S-6P-immunized RMs expressed high levels of Ki-67, confirming active proliferation. While most of these cells still expressed Ki-67 on day 14, the level of expression was strongly reduced, and by day 28, the majority of cells were Ki-67^-^ and had ceased to proliferate.

### B/HPIV3/S-6P immunization induces highly-functional SARS-CoV-2 S-specific Th1-biased CD4^+^ T-cells and cytotoxic CD8^+^ T-cells in the airways which transition to tissue-resident memory phenotypes

A more comprehensive functional analysis of the BAL-derived S-specific CD4^+^ T-cells revealed that, in addition to expressing IFNγ and TNFα, a proportion of these cells (about 40 to 80% from day 9 to 28 pi) also expressed IL-2, consistent with Th1 bias (Figure 5, A and B). Furthermore, a fraction of these S-specific CD4^+^ T-cells also expressed markers of cytotoxicity such as the degranulation markers CD107ab and granzyme B. Thus, the CD4^+^ T-cells induced by this vaccine displayed typical Th1-biased phenotype, similar to those generated after natural SARS-CoV-2 infection (DiPiazza et al., 2021; Jarjour et al., 2021; Szabo et al., 2021). S-specific CD8^+^ T-cells, in addition to expressing IFNγ and TNFα, also expressed high levels of degranulation markers CD107ab and granzyme B from day 9 to 28 pi, suggesting that these cells were highly functional (Figure 5, C and D). The phenotype of the blood-derived S-specific CD4^+^ and CD8^+^ T-cells was overall comparable to that of the airway-derived S-specific T-cells (Figure S5, A to D).

Furthermore, S-specific (IFNγ^+^/TNFα^+^) CD4^+^ and CD8^+^ T-cells from BAL could be separated into circulating CD69^-^ CD103^-^ and tissue-resident memory (Trm) CD69^+^ CD103^+/-^ subsets (Zheng and Wakim, 2021) (Figure 5, E and F for CD4^+^ T-cells and Figure 5, G and H for CD8^+^ T-cells, respectively). An additional subset of presumably tissue-resident S-specific CD4^+^ and CD8^+^ T-cells was identified as CD69^-^ CD103^+^ and has been previously detected in SARS-CoV-2-infected RMs (Nelson et al., 2022). Circulating CD69^-^ CD103^-^ S-specific CD4^+^ and CD8^+^ T-cells were detectable in BAL on day 9 pi and were prominent until day 14, representing about 60% of the S-specific T-cells on this day (Figure 5, F and H). Lung-resident S-specific CD69^+^ CD103^-^ CD4^+^ and CD8^+^ T-cells were detectable from day 9 pi (Figure 5, F and H), and their proportion increased through day 28 pi. On day 14, a fraction of these CD69^+^ S-specific T-cells had acquired CD103, and the proportion of CD69^+^ CD103^+^ Trm CD4^+^ and CD8^+^ T-cells further increased from day 14 to 28 pi. By day 28 pi, about 80% of the S-specific T cells in the airways were positive for CD103 and/or CD69, indicating transition of S antigen specific T cells to Trm phenotypes (Figure 5, F and H), while S-specific (IFNγ^+^/TNFα^+^) CD4^+^ and CD8^+^ T-cells in the blood mostly retained a circulating (CD69^-^ CD103^-^) phenotype throughout day 28 post-immunization (Figure S5, E to H).

Following antigen stimulation, the S-specific circulating and tissue-resident CD4^+^ or CD8^+^ T-cells recovered from the airways on days 9, 14, or 28 were phenotypically comparable with respect to strong expression of CD107ab and granzyme B (Figure S6), suggesting that all S specific CD69/CD103 subsets present in the airways were highly functional.

### B/HPIV3/S-6P immunization protects RM against SARS-CoV-2 challenge virus replication in the upper and lower airways

To assess protective efficacy of intranasal/intratracheal immunization with B/HPIV3/S-6P, we challenged RMs from both groups intranasally and intratracheally with 5.8 log_10_ TCID_50_ of SARS-CoV-2 WA1/2020 on day 30 or 31 after immunization (Figure S1). NS and BAL specimens were collected before challenge and on days 2, 4, and 6 post-challenge (pc). Viral RNA was extracted from these specimens, and the SARS-CoV-2 virus load was evaluated by RT-qPCR (Figure 6, A and B). We quantified subgenomic E (sgE) mRNA, indicative of SARS-CoV-2 infection, mRNA synthesis and challenge virus replication (Chandrashekar et al., 2020; Wolfel et al., 2020) (Figure 6, A and B, left panels). In B/HPIV3 empty-vector immunized RMs, sgE mRNA was detected in the UA of 4 of 4 and in the LA of 3 of 4 RMs, respectively, showing that all B/HPIV3 control immunized RM were actively infected with SARS-CoV-2 challenge virus. Copy numbers were maximal on day 2 pc (mean 5.0 log_10_ copies/ml in the UA, and 4.3 log_10_ copies/ml in the LA), and decreased until day 6 pc. In contrast, in all 4 B/HPIV3/S-6P-immunized RMs, sgE RNA was undetectable in the UA and LA at all time points (p<0.05), showing that intranasal/intratracheal immunization with a single dose of B/HPIV3/S-6P induces robust protection against high levels of challenge virus replication.

**Figure 6.**
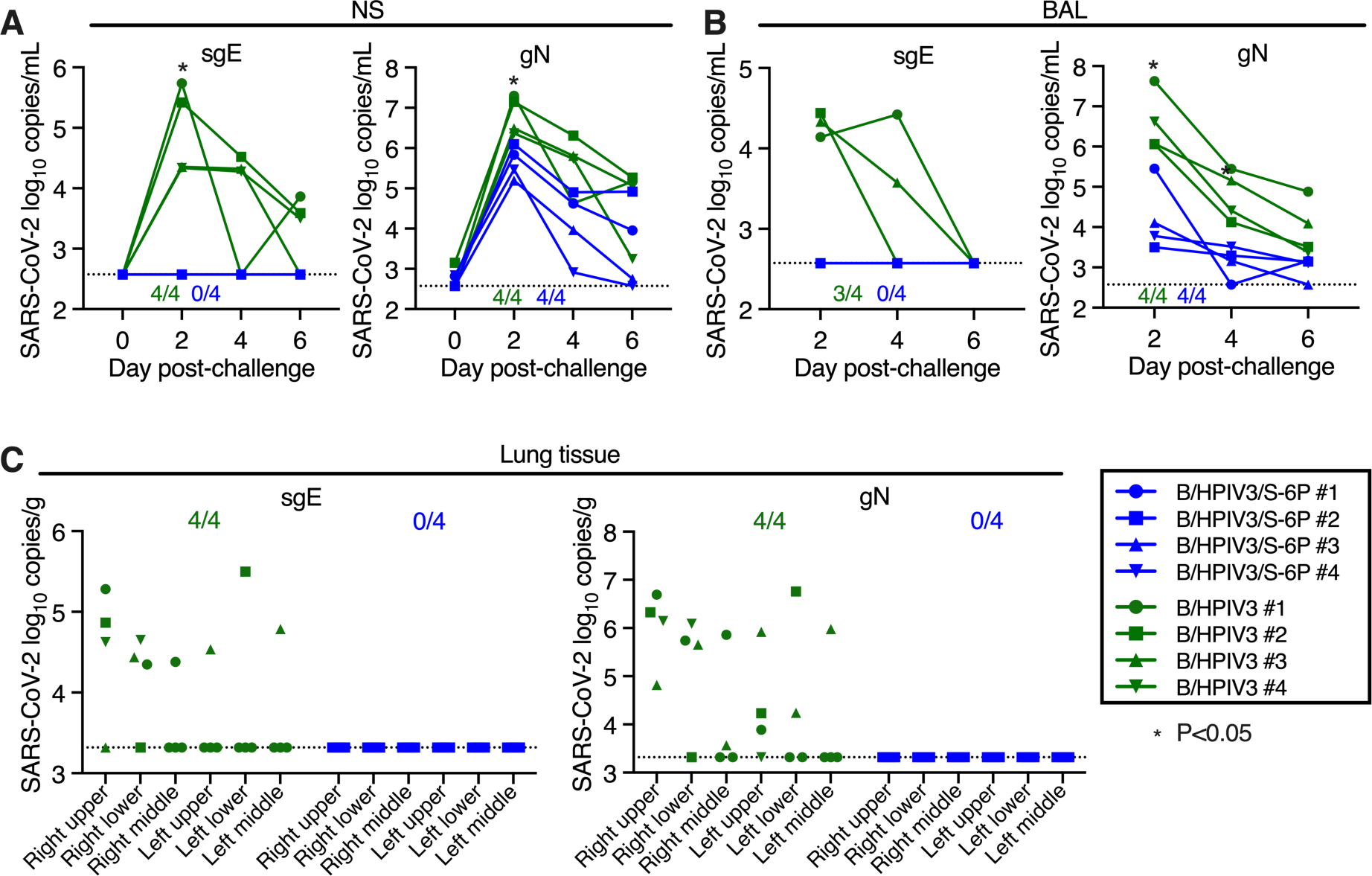
Absence of detectable SARS-CoV-2 challenge virus replication in the upper and lower airways and lung tissues of B/HPIV3/S-6P-immunized RMs. Rhesus macaques immunized with a single intranasal/intratracheal dose of B/HPIV3/S-6P or B/HPIV3 control (n=4 per group) were challenged intranasally/intratracheally on day 30 post-immunization with 5.8 TCID_50_ of SARS-CoV-2. (**A** and **B**) Evaluation of challenge virus shedding by qRT-PCR. (A) Nasal swabs (NS) and (B) bronchoalveolar lavage (BAL) fluid were collected on days 2, 4, and 6 post-challenge (pc), and SARS-CoV-2 subgenomic E mRNA (sgE, indicative of active SARS-CoV-2 replication) and genomic N RNA (gN, indicative of the presence of challenge virus) were quantified by RT-qPCR. (**C**) Challenge virus detection in lung tissues. Animals were euthanized on day 6 pc, and RNA was extracted from indicated areas of lung tissue for analysis by qRT-PCR. (A to C) The number of B/HPIV3/S-6P-immunized- or B/HPIV3-immunized RMs with sgE or gN RNA detectable by RT-qPCR in each set of samples is indicated. The limit of detection was 2.6 log_10_ copies per ml of NS or BAL fluid and 3.3 log_10_ copies per g of lung tissue. B/HPIV3/S-6P-immunized RMs are shown in blue, B/HPIV3-immunized RMs are in green, with each RM indicated by a symbol. *P<0.05.

From the same samples, we also quantified genomic N (gN), indicative of the presence of challenge virus. gN RNA copies/ml were maximal on day 2 pi in the UA and LA of RMs, and then steadily decreased over time. The average of the gN RNA load on day 2 in the UA of RMs immunized with B/HPIV3/S-6P was 16-fold lower than that of RMs immunized with B/HPIV3 empty vector control (6.8 log_10_ vs 5.6 log_10_ copies/ml in the B/HPIV3- and the B/HPIV3/S-6P-immunized RMs, respectively, P<0.05). In the LA, the average of the genomic N RNA load in B/HPIV3/S-6P-immunized RMs was 240-fold lower than in RMs immunized with B/HPIV3 (6.6 log_10_ vs 4.2 log_10_ copies/mL in the B/HPIV3- and the B/HPIV3/S-6P-immunized RMs, respectively, P<0.05), which would be consistent with the presence of residual challenge virus, and absence of SARS-CoV-2 challenge virus replication, in B/HPIV3/S-6P immunized animals.

Quantification of sgE and gN RNA was also performed for lung tissues from different areas obtained on day 6 pc (Figure 6C). In all 4 B/HPIV3 empty-vector immunized RMs, gN RNA and sgE mRNA were detected, mostly in the right upper and right lower lobes of the lungs. Neither gN RNA nor sgE mRNA were detected in the lungs of the B/HPIV3/S-6P-immunized RMs, confirming robust protection against SARS-CoV-2 infection induced by B/HPIV3/S-6P. Furthermore, no active SARS-CoV-2 replication was detected from rectal swab samples (Figure S8).

We finally assessed the CD4^+^ and CD8^+^ T-cell response in the blood (Figure 4, B and C) and lower airways (Figure 4, E and F) of immunized RMs, 4 days after challenge with SARS-CoV-2. In the blood, an increase of S-specific IFNγ^+^/TNFα^+^ CD4^+^ and CD8^+^ T-cells was detected in 3 and 1 of 4 B/HPIV3/S-6P-immunized RMs, respectively, that correlated well with the increased expression of Ki-67 by the S-specific CD4^+^ T-cells (Figure S7C). A modest increase of S-specific IFNγ^+^/TNFα^+^ CD4^+^ T-cells was also detected in 1 B/HPIV3-immunized RM (Figure 4B). However, in the lower airways, a decrease rather than an increase of the S-specific IFNγ^+^/TNFα^+^ CD4^+^ and CD8^+^ T-cells was detected in the B/HPIV3/S6-P-immunized RMs and no active T-cell proliferation was detected (Figure S7D).

## DISCUSSION

SARS-CoV-2 vaccines for infants and young children are critically needed. Equally needed are topical SARS-CoV-2 vaccines that directly stimulate local respiratory tract immunity in addition to systemic immunity, which might effectively reduce infection and transmission. In the present study, we evaluated a chimeric B/HPIV3 virus as a live topical viral vector to express the SARS-CoV-2 S protein, stabilized in its prefusion form, expected to be non-functional for virus entry (Hsieh et al., 2020). No effects of B/HPIV3 and B/HPIV3/S-6P on the general health of rhesus macaques were observed following immunization, indicating that this vector and the expressed S protein were safe in this non-human primate model.

The B/HPIV3 vector is in clinical development and is predicted to be safe. The HN and F surface glycoproteins of B/HPIV3 are derived from HPIV3. HN and F are the major protective antigens of HPIV3, a major respiratory pathogen in infants and children under 5 years of age. The BPIV3 genes encode for the nucleocapsid N, phosphoprotein P, interferon antagonist V, matrix M and polymerase L that provide host range restriction and represent the basis for the strong attenuation of this vector while the surface glycoproteins from HPIV3 mediate infection (Bernstein et al., 2012; Karron et al., 2012). The B/HPIV3 vector exhibits a natural tropism for the respiratory tract without any evidence of spread to other tissues. An important advantage of the B/HPIV3 vector is that it has been evaluated in young children both as the empty vector and expressing a foreign gene (RSV F protein), and in both cases it had an excellent clinical safety profile (Bernstein et al., 2012; Karron et al., 2012). This should expedite advancing this SARS-CoV-2 vaccine through clinical studies. A second important advantage of the B/HPIV3 vector is that it is administered intranasally and stimulates systemic and local mucosal immunity in the respiratory tract, the site of SARS-CoV-2 entry, replication, disease, and egress. Finally, another bonus associated with the use of B/HPIV3 as a vector for expressing a heterologous viral protein is that it also induces neutralizing antibody responses against HPIV3 itself (Bernstein et al., 2012; Karron et al., 2012), effectively serving as a dual HPIV3/SARS-CoV-2 vaccine.

A single intranasal/intratracheal immunization with B/HPIV3/S-6P efficiently induced mucosal IgA and IgG in the UA and LA of all immunized RMs, as well as strong serum IgM, IgA and IgG responses to SARS-CoV-2 S protein and its RBD. The anti-S and anti-RBD IgG responses were comparable to those detected in human convalescent plasma of individuals with high levels of anti-S and anti-RBD IgG antibodies. The serum antibodies efficiently neutralized the vaccine-matched SARS-CoV-2 WA1/2020 strain, as well as VoCs of B.1.1.7/Alpha and B.1.617.2/Delta lineages. However, the neutralizing activity of these sera against the B.1351/Beta and B.1.1.529/Omicron lineages was limited. This suggests that a boost to a primary immunization with B/HPIV3/S-6P might be required, as with perhaps any SARS-CoV-2 vaccine, to boost antibody titers and affinity maturation, providing for greater breadth of antigen recognition and protection against VoCs.

While not specifically investigated, B/HPIV3/S-6P immunization likely resulted in the generation of long-lived antigen-specific tissue-resident B and T-cells in the mucosa. Antigen-specific B cells in the mucosa are sources of soluble dimeric IgA (Oh et al., 2021) with increased neutralizing activity compared to serum-derived monomeric IgA (Wang et al., 2021). Injectable SARS-CoV-2 vaccines have been shown to result in the appearance of mucosal anti-S IgA and IgG in the URT and LRT of immunized RM, primarily by transudation of monomeric Igs from the blood, and in absence of direct stimulation of mucosal immunity at the site of infection (Corbett et al., 2021a). To induce long-term mucosal humoral immunity in the airways, a homologous or heterologous prime/boost approach that includes an intranasal vaccine might be advantageous over a prime/boost approach based solely on immunizations with injectable vaccines.

B/HPIV3/S-6P also induced S-specific CD4^+^ and CD8^+^ T-cells in the blood and the LA. Similar to immunization with injectable SARS-CoV-2 vaccines, intranasal/intratracheal immunization with B/HPIV3/S-6P induced S-specific Th1-biased CD4^+^ T-cells in the blood that expressed IFNβ, TNFα, and IL-2 (Corbett et al., 2020b; Corbett et al., 2021a; Corbett et al., 2021b; Joyce et al., 2022). Furthermore, these Th1-biased S-specific CD4^+^ T-cells expressed markers of cytotoxicity such as CD107ab and granzyme B, suggesting that they might also be directly involved in virus clearance. In addition, B/HPIV3/S-6P appeared to induce a stronger response of S-specific CD8^+^ T-cells in the blood of RMs than injectable vaccines (Corbett et al., 2020b; Corbett et al., 2021b; Mercado et al., 2020).

To control respiratory virus infections, mucosal immune responses at the site of infection are essential (Sette and Crotty, 2021). In longitudinal studies of SARS-CoV-2 infection in humans, T-cells induced in airways exhibited activated and tissue-resident signatures and functionally protective profiles (Poon et al., 2021), and their frequencies in airways (but not in blood) correlated with younger age and survival (Rydyznski Moderbacher et al., 2020; Szabo et al., 2021). Depletion of CD8^+^ T-cells partially abrogated the protective immunity against SARS-CoV-2 re-challenge in the macaque model (McMahan et al., 2021). Intranasal immunization with B/HPIV3/S-6P induced strong T-cell responses in the LA, with a very efficient induction of S-specific Th1-biased CD4^+^ T-cells expressing IFNβ, TNFα, and IL-2 as well as S-specific CD8^+^ T-cells that expressed IFNβ, TNFα, and granzyme B, while injectable mRNA or adenovirus-based vaccines do not appear to induce S-specific CD4^+^ and CD8^+^ T-cells in the airways of immunized RMs (Corbett et al., 2020b; Corbett et al., 2021b). A substantial fraction of the S-specific CD4^+^ and CD8^+^ T-cells in the lungs expressed the Trm marker CD69, with a subpopulation also expressing CD103. The proportion of S-specific Trm T-cells increased over time and represented the main S-specific T-cell population one month after immunization. While the contribution of memory CD4^+^ and CD8^+^ T-cells in controlling SARS-CoV-2 replication is still not fully understood (Hasenkrug et al., 2021; McMahan et al., 2021; Nomura et al., 2021; Tan et al., 2021), previous work suggested that Trm T-cells contribute to long-term immunity and are associated with protection against respiratory disease (Auladell et al., 2019; Kohlmeier et al., 2007; Szabo et al., 2021; Tan et al., 2021). Furthermore, as T-cell epitopes in the S protein are mostly conserved across SARS-CoV-2 lineages (Keeton et al., 2022; Tarke et al., 2021), the strong induction of S-specific Trm CD4^+^ and CD8^+^ T-cells by B/HPIV3/S-6P should be beneficial to the protection against VoCs including Omicron (Auladell et al., 2019; Jarjour et al., 2021; Sette and Crotty, 2021). Indeed, in the rhesus model, the failure of injectable SARS-CoV-2 vaccines in controlling SARS-CoV-2 B.1.529/Omicron challenge virus replication in the upper respiratory tract was associated with negligible T-cell responses, but not with the level of anti-S antibody titers (Chandrashekar et al., 2022). In addition, even though the S expression by B/HPIV3/S-6P in the LA of one RM was not detected (Figure S2), this RM still exhibited strong serum and mucosal anti-S IgA and IgG response as well as strong S-specific CD4^+^ and CD8^+^ T-cells in the blood and LA. This suggested that even a low level of S expression in the airways was highly immunogenic.

We found that RMs were fully protected from SARS-CoV-2 challenge 1 month after immunization. No SARS-CoV-2 challenge virus replication was detectable in the UA or LA or in lung tissues of immunized RMs, suggesting sterilizing immunity under these experimental conditions. However, the duration of immunity is currently unknown and will be evaluated in a separate study. Following challenge, we detected a recall response of S-specific CD4^+^ T-cells in the blood of B/HPIV3/S-6P-immunized RMs, but not in the LA. The reasons for the apparent absence of a detectable recall response are still unknown. It is possible that the S-specific T-cells were not present in the airways and inaccessible through BAL, or that the BAL cell collection on day 4 pc was too early to detect any recall response, or that in absence of challenge virus replication, a recall response was not triggered in the LA. Later time points following challenge will be included in future studies. Even though we did not evaluate protection against HPIV3 challenge in this study, we showed that B/HPIV3 induced strong serum HPIV3 neutralizing antibodies, comparable to levels seen in previous studies (Schmidt et al., 2001); thus, B/HPIV3/S-6P represents a promising candidate for intranasal immunization against both SARS-CoV-2 and HPIV3, an important pediatric cause of lower respiratory tract disease.

In conclusion, a single topical immunization with B/HPIV3/S-6P was highly immunogenic and protective against SARS-CoV-2 in RMs. Our data support the further development of this vaccine candidate for potential use as a stand-alone vaccine and/or in a prime/boost combination with other SARS-CoV-2 vaccines for infants and young children. The B/HPIV3/S-6P vaccine candidate will be evaluated in a Phase 1 study.

## ACKNOWLEDGEMENTS

We thank Alicia Wojcik and Amanda Havenner and the staff of the National Institute of Allergy and Infectious Diseases (NIAID) Comparative Medicine Branch for animal study support, Jeffrey I. Cohen for providing plasma from SARS-CoV-2 convalescent individuals, and Peter L. Collins for helpful discussions and for comments on the manuscript.

## AUTHOR CONTRIBUTIONS

Conceptualization: CLN, DB, UJB Design of Experiments: CLN, CEN, XL, HSP, YM, CL, RH, AC, RM, PL, SM, DB, UJB Investigation: CLN, CEN, XL, HSP, YM, CL, CS, LY, INM, TWK, AW, RM, PZ, LEV

Reagent development: RFJ, NLG

Data analysis and visualization: CLN, CEN, XL, HSP, YM, CL, INM, PZ, PL, DB, UJB Writing – original draft: CLN, DB, UJB

Writing – review and editing: CLN, CEN, XL, HSP, YM, CL, CS, LY, RH, AC, INM, TWK, AW, RM, PZ, PL, RFJ, NLG, LEV, SM, DB, UJB

## STAR METHODS

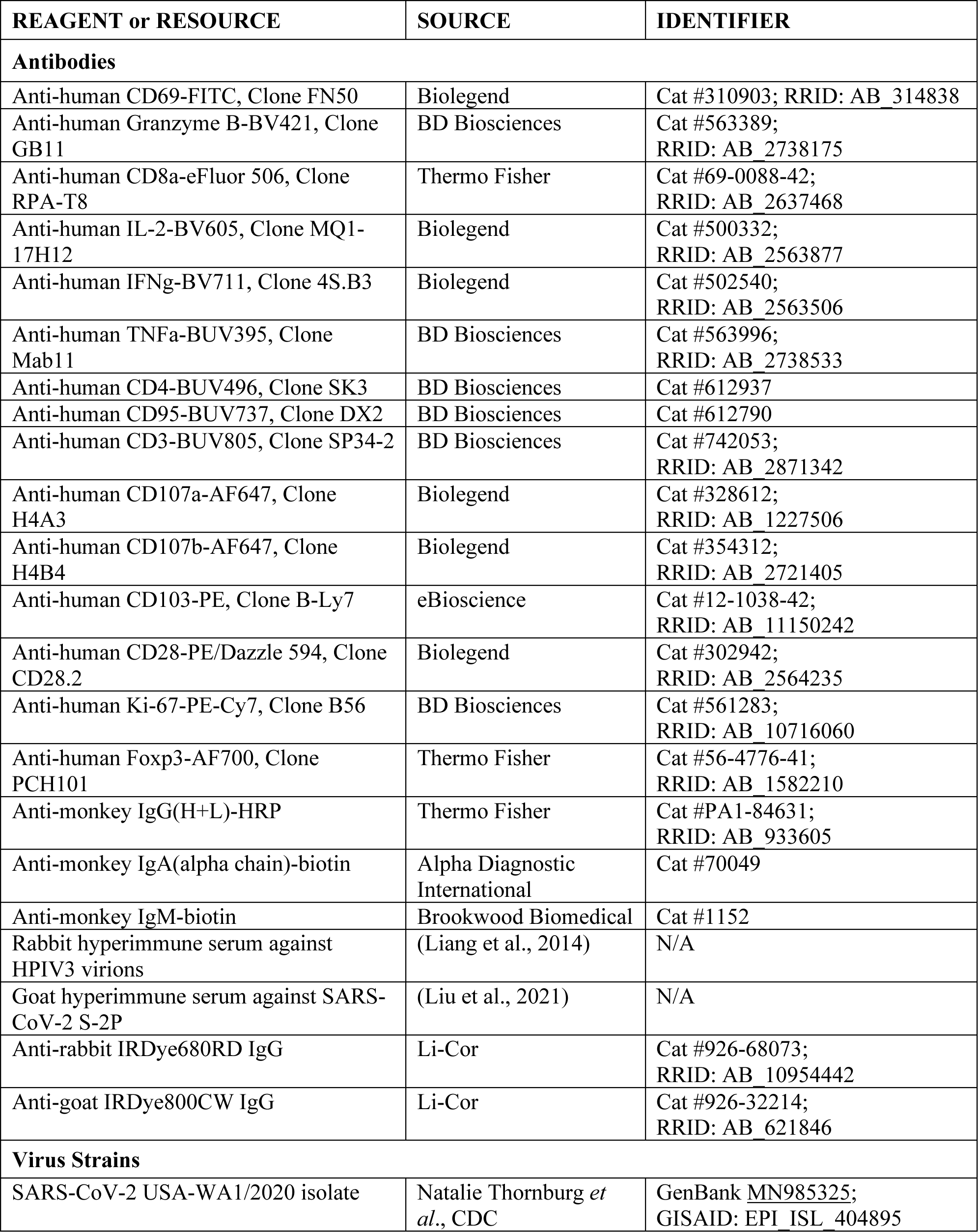
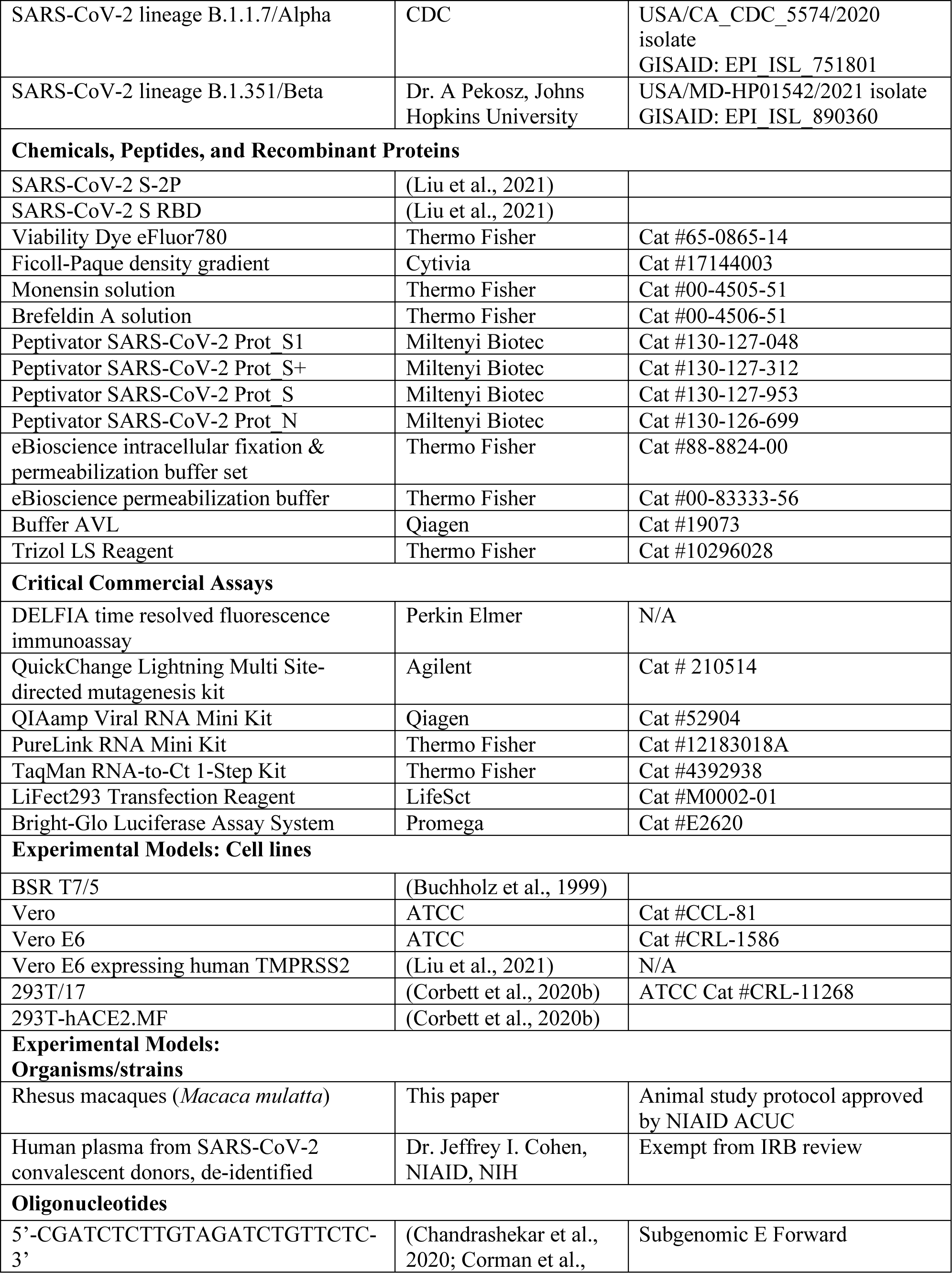
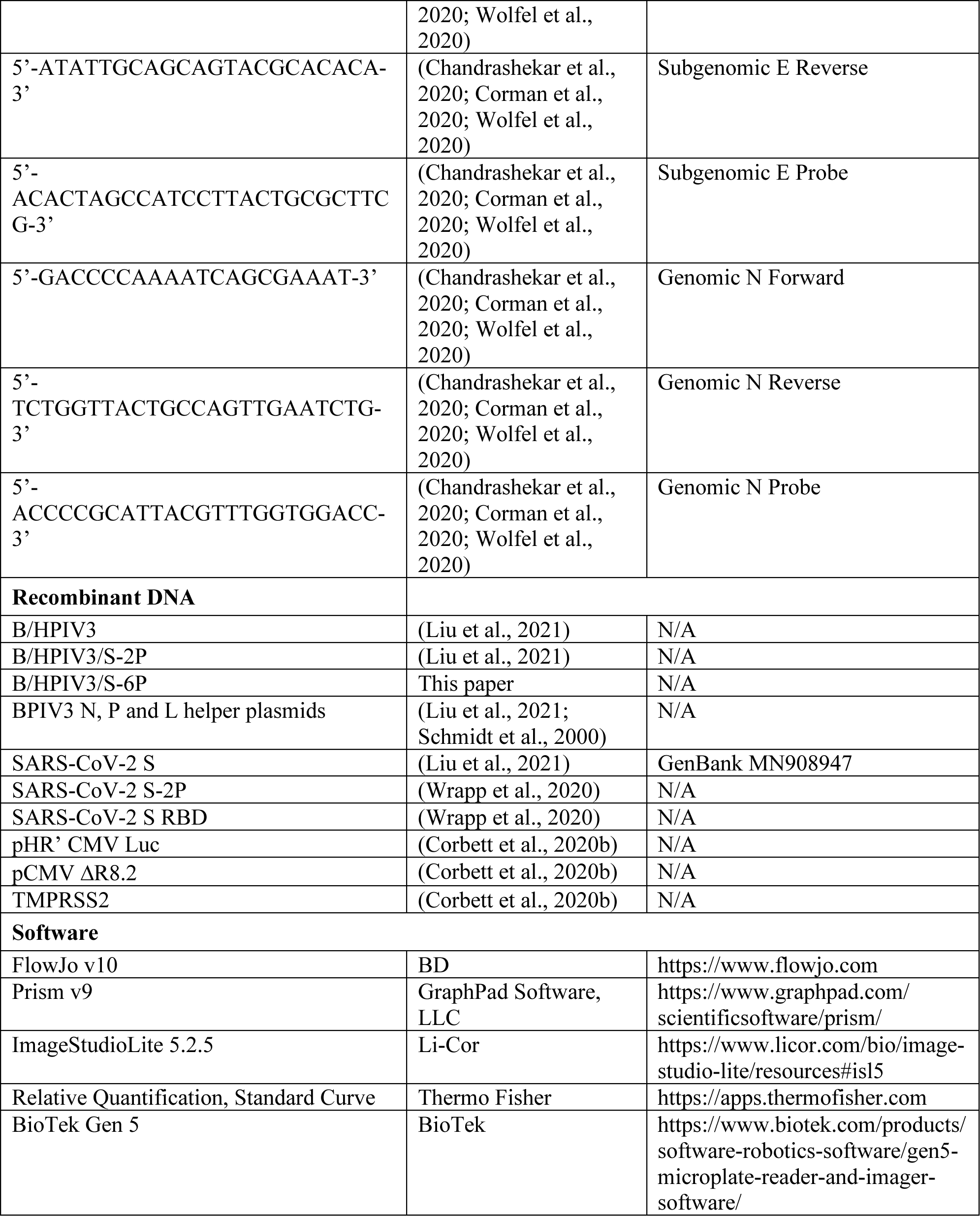

### RESOURCE AVAILABILITY

Requests for resources, reagents and further information regarding this manuscript should be addressed and fulfilled by the lead contact, Ursula Buchholz.

#### Materials availability

Plasmids and viruses newly generated in this study are available under material transfer request upon request to the lead contact. All data are included in the manuscript.

### EXPERIMENTAL MODEL DETAILS

#### Cell lines

Baby hamster kidney cells expressing T7 RNA polymerase (BSR T7/5) were grown in Glasgow minimum essential medium (MEM) (Thermo Fisher Scientific) with 10% Fetal bovine serum (FBS), 1% L-glutamine (Thermo Fisher Scientific), and 2% MEM Amino Acids (Thermo Fisher Scientific). African green monkey kidney Vero (ATCC CCL-81) and Vero E6 (ATCC CRL-1586) cells were cultured in Dulbecco’s MEM with GlutaMAX (Thermo Fisher Scientific) with 5% FBS and 1% L-glutamine. Vero E6 cells that express high levels of ACE2 (Chu et al., 2020; Ren et al., 2006) and Vero E6 cells that stably express human TMPRSS2 (Liu et al., 2021) were used to expand SARS-CoV-2.

#### SARS-CoV-2 virus stocks

The SARS-CoV-2 USA-WA1/2020 isolate (lineage A; GenBank <u>MN985325</u>; GISAID: EPI_ISL_404895; obtained from Dr. Natalie Thornburg *et al*., Centers for Disease Control and Prevention (CDC)) was passaged on Vero E6 cells. We used WA1/2020 as challenge virus in this study because its S sequence is homologous to the sequence from which S-6P was derived. The USA/CA_CDC_5574/2020 isolate [lineage B.1.1.7 (Alpha); GISAID: EPI_ISL_751801; obtained from CDC] and the USA/MD-HP01542/2021 isolate [lineage B.1.351 (Beta); GISAID: EPI_ISL_890360; obtained from Dr. Andrew Pekosz, Johns Hopkins University] were grown on TMPRSS2-expressing Vero E6 cells (Liu et al., 2021). The SARS-CoV-2 stocks were titrated in Vero E6 cells by determination of the 50% tissue culture infectious dose (TCID_50_)(Subbarao et al., 2004). All experiments with SARS-CoV-2 were performed in biosafety level 3 (BSL3) containment laboratories approved by the USDA and CDC.

### METHOD DETAILS

#### Generation of the B/HPIV3/S-6P vaccine

The B/HPIV3/S-6P vaccine candidate is an improved derivative of B/HPIV3/S-2P (Liu et al., 2021). To generate the B/HPIV3/S-2P cDNA (Liu et al., 2021), the ORF encoding the full-length 1,273 aa SARS-CoV-2 S protein from the first available sequence (GenBank MN908947, Wuhan-Hu-1; amino acid sequence of the S protein identical to that of WA1/2020) was codon-optimized for human expression and synthesized commercially (BioBasic). Two proline substitutions (aa positions 986 and 987) and four aa substitutions (RRAR to GSAS, aa 682-685) that stabilize S in the prefusion conformation and ablate the furin cleavage site between S1 and S2 (Wrapp et al., 2020) were introduced to generate the S-2P cDNA (Liu et al., 2021). This S-2P ORF was then inserted into a cDNA clone encoding the B/HPIV3 antigenome between the N and P ORFs to create the B/HPIV3/S-2P cDNA (Liu et al., 2021). To create the B/HPIV3/S-6P cDNA, the B/HPIV3/S-2P cDNA was modified to introduce 4 additional proline substitutions in the S ORF (aa position 817, 892, 899, and 942 for a total of 6 proline substitutions). The 4 additional proline substitutions had been shown to confer increased stability to a soluble version of the prefusion-stabilized S protein (Hsieh et al., 2020). The B/HPIV3/S-6P cDNA was used to transfect BHK21 cells (clone BSR T7/5, stably expressing T7 RNA polymerase (Buchholz et al., 1999)), together with helper plasmids encoding the N, P and L proteins (Buchholz et al., 2004; Liu et al., 2021), to produce the B/HPIV3/S-6P recombinant virus. The empty vector control virus B/HPIV3 was recovered in parallel. Virus stocks were grown in Vero cells, and viral genomes of recovered viruses were completely sequenced by Sanger sequencing using overlapping RT-PCR fragments, confirming the absence of any adventitious mutations.

#### Immunization and challenge of rhesus macaques

All animal studies were approved by the NIAID Animal Care and Use Committee. The timeline of the experiment and sampling is summarized in Figure S1. Eight juvenile to young adult male Indian-origin rhesus macaques (*Macaca mulatta*), confirmed to be seronegative for HPIV3 and SARS-CoV-2, were immunized intranasally (0.5 ml per nostril) and intratracheally (1 ml) with a total does of 6.3 log_10_ plaque-forming units (PFU) of B/HPIV3/S-6P or the empty vector control B/HPIV3. Animals were observed daily from day -3 until the end of the study. Each time they were sedated, animals were weighed, their rectal temperature was taken, as well as the pulse in beats per minute and the respiratory rate in breaths per minute. In addition, the blood oxygen levels were determined by pulse oximetry.

Blood for analysis of serum antibodies and peripheral blood mononuclear cells (PBMC) was collected on days -3, 4, 9, 14, 21, and 28 pi. Nasopharyngeal swabs (NS) for vaccine virus quantification in the upper airways (UA) were performed daily from day -3 to day 10 pi and on days 12 and 14 pi using cotton-tipped applicators. Swabs were placed in 2 ml Leibovitz (L-15) medium with 1x sucrose phosphate (SP) used as stabilizer, and vortexed for 10 seconds. Aliquots were then snap frozen in dry ice and stored at -80°C. Nasal washes (NWs) for analysis of mucosal IgA and IgG were performed using 1 ml of Lactated Ringer’s solution per nostril (2 ml total) on days -3, 14, 21 and 28 pi and aliquots were snap frozen in dry ice and stored at -80°C until further analysis. Tracheal lavages (TL) for virus quantification were done every other day from day 2 to 8 pi and on day 12 pi using 3 ml PBS. The samples were mixed 1:1 with L-15 medium containing 2x SP and aliquots were snap frozen in dry ice and stored at -80°C for further analysis. Bronchoalveolar lavages (BALs) for analysis of mucosal IgA and IgG and airway immune cells from the lower airways (LA) were done on days -3, 9, 14 and 28 pi using 30 ml PBS (3 times 10 ml). For analysis of mononuclear cells, BAL was filtered through a 100 µm filter, and centrifuged at 544 x g for 15 min at 4°C. The cell pellet was resuspended at 2x10^7^ cells/ml in X-VIVO 15 media supplemented with 10% FBS for subsequent analysis. The cell-free BAL was aliquoted, snap frozen in dry ice and stored at -80°C for further analysis. Rectal swabs were done on day -3 and then every other day from day 2 to 14 following the same procedure as NS.

Four weeks after immunization, animals were transferred to BSL3. On day 30 or 31 pi, animals were challenged intranasally and intratracheally on with 10^5.8^ TCID_50_ of SARS-CoV-2, USA-WA1/2020, that was entirely sequenced and free of any prominent adventitious mutations. Sample collections were done following the same procedures as during the immunization phase. Briefly, blood was collected before challenge and on day 6 post-challenge (pc). NS were performed every other day from day 0 to day 6 pc. NWs were done on day 6 pc, BAL on days 2, 4 and 6 pc, and rectal swabs on days 0, 2, 4 and 6 pc. Animals were necropsied on day 6 pc, and tissues were collected. 6 samples per animal from individual lung lobes were collected, and snap frozen in dry ice for further analysis.

#### Immunoplaque assay for titration of B/HPIV3 and B/HPIV3/S-6P

Titers of B/HPIV3 and B/HPIV3/S-6P from NS and TLs were determined by dual-staining immunoplaque assay (Liu et al., 2021). Briefly, Vero cell monolayers in 24-well plates were infected in duplicate with 10-fold serially diluted samples. Infected monolayers were overlaid with culture medium containing 0.8% methylcellulose, and incubated at 32°C for 6 days, fixed with 80% methanol, and immunostained with a rabbit hyperimmune serum raised against purified HPIV3 virions to detect B/HPIV3 antigens, and a goat hyperimmune serum to the secreted SARS-CoV-2 S to detect co-expression of the S protein, followed by infrared-dye conjugated donkey anti-rabbit IRDye680 IgG and donkey anti-goat IRDye800 IgG secondary antibodies (LiCor). Plates were scanned with the Odyssey infrared imaging system (LiCor).

Fluorescent staining for PIV3 proteins and SARS-CoV-2 S was visualized in green and red, respectively, providing for yellow plaque staining when merged.

#### Dissociation-enhanced lanthanide fluorescent (DELFIA) time resolved fluorescence (TRF) immunoassay, ELISA and live HPIV3 and SARS-CoV-2 neutralization assay

Levels of anti-SARS-CoV-2 S antibodies elicited by B/HPIV3/S-6P were determined by DELFIA-TRF (Perkin Elmer) from NW or BAL following the supplier’s protocol and from serum samples by ELISA (Liu et al., 2021) using the recombinantly-expressed secreted version of S-2P (Wrapp et al., 2020), or a fragment (aa 328-531) containing the receptor binding domain (RBD) of the SARS-CoV-2 S protein (Walls et al., 2020). The secondary antibodies used in both assays were goat anti-monkey IgG(H+L) horseradish peroxidase (HRP) (Thermo Fisher, Cat #PA1-84631), goat anti-monkey IgA (alpha chain)-biotin (Alpha Diagnostic International, Cat #70049), and goat anti-monkey IgM-biotin (Brookwood Biomedical, Cat#1152).

HPIV3-specific neutralizing antibody titers were measured by a 60% plaque reduction neutralization test (PRNT_60_) (Liu et al., 2021). The serum neutralizing antibody assays using live SARS-CoV-2 virus was performed in a BSL3 laboratory. Heat-inactivated sera were 2-fold serially diluted in Opti-MEM and mixed with an equal volume of SARS-CoV-2 (100 TCID_50_) and incubated at 37°C for 1 h. Mixtures were added to quadruplicate wells of Vero E6 cells in 96-well plates and incubated for four days. The 50% neutralizing dose (ND_50_) was defined as the highest dilution of serum that completely prevented cytopathic effect in 50% of the wells and was expressed as a log_10_ reciprocal value (Liu et al., 2021).

#### Lentivirus-based pseudotype virus neutralization assay

The SARS-CoV-2 pseudovirus neutralization assays were performed as previously reported (Corbett et al., 2020b). Briefly, the single-round luciferase-expressing pseudoviruses were generated by co-transfection of plasmids encoding SARS-CoV-2 S of isolate Wuhan-Hu-1, GenBank accession number MN908947.3, or of lineages B.1.351/Beta, B.1.1.7/Alpha, B.1.617.2/Delta, B.1.1.529/Omicron), luciferase reporter (pHR’ CMV Luc), lentivirus backbone (pCMV ΔR8.2), and human transmembrane protease serine 2 (TMPRSS2) at a ratio of 1:20:20:0.3 into HEK293T/17 cells (ATCC) with transfection reagent LiFect293™. The pseudoviruses were harvested at 72 h post transfection. The supernatants were collected after centrifugation at 478 x g for 10 minutes to remove cell debris, then filtered through a 0.45 mm filter, aliquoted and titrated before neutralization assay. For the antibody neutralization assay, 6-point, 5-fold dilution series were prepared in culture medium (DMEM medium with 10% FBS, 1 % Pen/Strep and 3 µg/ml puromycin). Fifty µl antibody dilution were mixed with 50 µl of diluted pseudoviruses in the 96-well plate and incubated for 30 min at 37°C. Ten thousand ACE2-expressing 293T-cells (293T-hACE2.MF stable cell line cells) were added in a final volume of 200 µl. Seventy-two h later, after carefully removing all the supernatants, cells were lysed with Bright-Glo™ luciferase assay substrate (Promega), and luciferase activity (relative light units, RLU) was measured. Percent neutralization was normalized relative to uninfected cells as 100% neutralization and cells infected with only pseudoviruses as 0% neutralization. IC_50_ titers were determined using a log (agonist) vs. normalized response (variable slope) nonlinear function in Prism v8 (GraphPad).

#### Evaluation of the T cell response in the blood and lower airways of RMs

Blood and BAL collection procedures followed ACUC-approved standard operating procedures and limits. Blood that was collected in EDTA tubes was diluted 1:1 with PBS. Fifteen ml of Ficoll-Paque density gradient (GE Healthcare) was added to Leucosep PBMC isolation tubes (Greiner bio-one) and centrifuged at 1,000 x g for 1 min at 22°C to collect Ficoll below the separation filter. The blood and PBS mixture was added to the Leucosep tubes with Ficoll-Paque and centrifuged at 863 x g for 10 min at 22°C. The upper layer was poured into a 50 ml conical tube and brought to 50 ml with PBS, and then centrifuged at 544 x g for 5 min at 4°C. The cell pellet was resuspended at 2x10^7^ cell/ml in 90% FBS and 10% DMSO for storage at -80°C overnight before being transferred into liquid nitrogen.

Single cell suspensions of PBMCs that had been rested overnight or freshly collected BAL cells were plated at 2x10^7^ cells/ml in 200 ml in 96 well plates with X-VIVO 15 media, with 10% FBS, 1000x Brefeldin (Thermo Fisher Cat#00-4506-51) and 1000x Monensin (Thermo Fisher Cat#00-4505-51), CD107a APC 1:50, CD107b APC 1:50, and the indicated peptide pools at 1mg/ml. Replica wells were not stimulated. Spike peptide pools consisted of Peptivator SARS-CoV-2 Prot_S1 (Miltenyi Cat#130-127-048), Peptivator SARS-CoV-2 Prot_S+ (Miltenyi Cat# 130-127-312), and Peptivator SARS-CoV-2 Prot_S (Miltenyi Cat#130-127-953) covering the whole spike protein. Nucleocapsid peptide pool consisted of Peptivator SARS-CoV-2 Prot_N (Miltenyi Cat# 130-126-699). Cells were stimulated for 6 h at 37°C with 5% CO_2_. After stimulation, cells were centrifuged at 544 x g for 5 min at 4°C, and further processed by surface staining.

Cells were resuspended in 50 ml surface stain antibodies diluted in PBS with 1% FBS and incubated for 20 min at 4°C. Cells were washed 3 times with PBS with 1% FBS before fixation with intracellular fixation & permeabilization buffer set (Thermo Fisher, Cat# 88-8824-00) for 16 h at 4°C. After fixation, cells were centrifuged at 1,028 x g for 5 min at 4°C without brake and washed once with permeabilization buffer. Cells were resuspended in 50 ml intracellular stains diluted in permeabilization buffer, and stained for 30 min at 4°C. The antibodies used for extracellular and intracellular staining were: CD69 (FITC, clone FN50, Biolegend), granzyme B (BV421, clone GB11, BD Biosciences), CD8a (eFluor 506, clone RPA-T8, Thermo Fisher), IL-2 (BV605, MQ-17H12, Biolegend), IFNγ(BV711, clone 4S.B3, Biolegend), IL-17 (BV785, clone BL168, Biolegend), TNFα (BUV395, clone Mab11, BD Biosciences), CD4 (BUV496, clone SK3, BD Biosciences), CD95 (BUV737, clone DX2, BD Biosciences), CD3 (BUV805, clone SP34-2, BD Biosciences), CD107a (AF647, clone H4A3, Biolegend), CD107b (AF647, clone H4B4, Biolegend), viability Dye eFluor780 (Thermo Fisher), CD103 (PE, clone B-Ly7, eBioscience), CD28 (PE/Dazzle 594, clone CD28.2, Biolegend), Ki-67 (PE-Cy7, clone B56, BD Biosciences), Foxp3 (AF700, clone PCH101, Thermo Fisher). After staining, cells were washed with 2x eBioscience permeabilization buffer 2x and resuspended in PBS supplemented with 1% FBS and 0.05% sodium azide for flow cytometry analysis on the BD Symphony platform. Data were analyzed using FlowJo version 10.

#### Quantification of SARS-CoV-2 genomic and subgenomic RNA

Hundred ml each of NS and BAL fluid collected on day 2, 4 and 6 pc and rectal swabs collected on day 6 pc were inactivated in a BSL3 laboratory using 400 ml buffer AVL (Qiagen) and 500 ml ethanol, and RNA was extracted using the QIAamp Viral RNA Mini Kit (Qiagen) according to the manufacturer’s protocol. To extract total RNA from lung homogenates harvested on day 6 pc, 300 ml of each lung homogenate (at a concentration of 0.1 g of tissue/ml) was mixed with 900 ml TRIzol LS (Thermo Fisher) using Phasemaker Tubes (Thermo Fisher) and RNA was extracted using the PureLink RNA Mini Kit (Thermo Fisher) following the manufacturer’s instructions. Then, the SARS-CoV-2 genomic N RNA and subgenomic E mRNA were quantified in triplicate using the TaqMan RNA-to-Ct 1-Step Kit (Thermo Fisher) using previously reported TaqMan primers/probes (Chandrashekar et al., 2020; Corman et al., 2020; Wolfel et al., 2020) on the QuantStudio 7 Pro (ThermoFisher). Standard curves were generated using serially diluted pcDNA3.1 plasmids encoding gN, gE, or sgE sequences. The limit of detection was 2.57 log_10_ copies per ml of NS, BAL fluid, or rectal swabs and 3.32 log_10_ copies per g of lung tissue.

#### Statistical analysis

Data sets were assessed for significance using two-way ANOVA with Sidak’s multiple comparison test using Prism 8 (GraphPad Software). Data were only considered significant at P < 0.05.

