## Supplemental Figures S1-S8 for "Intranasal pediatric parainfluenza virus-vectored SARS-CoV-2 vaccine candidate is protective in macaques"

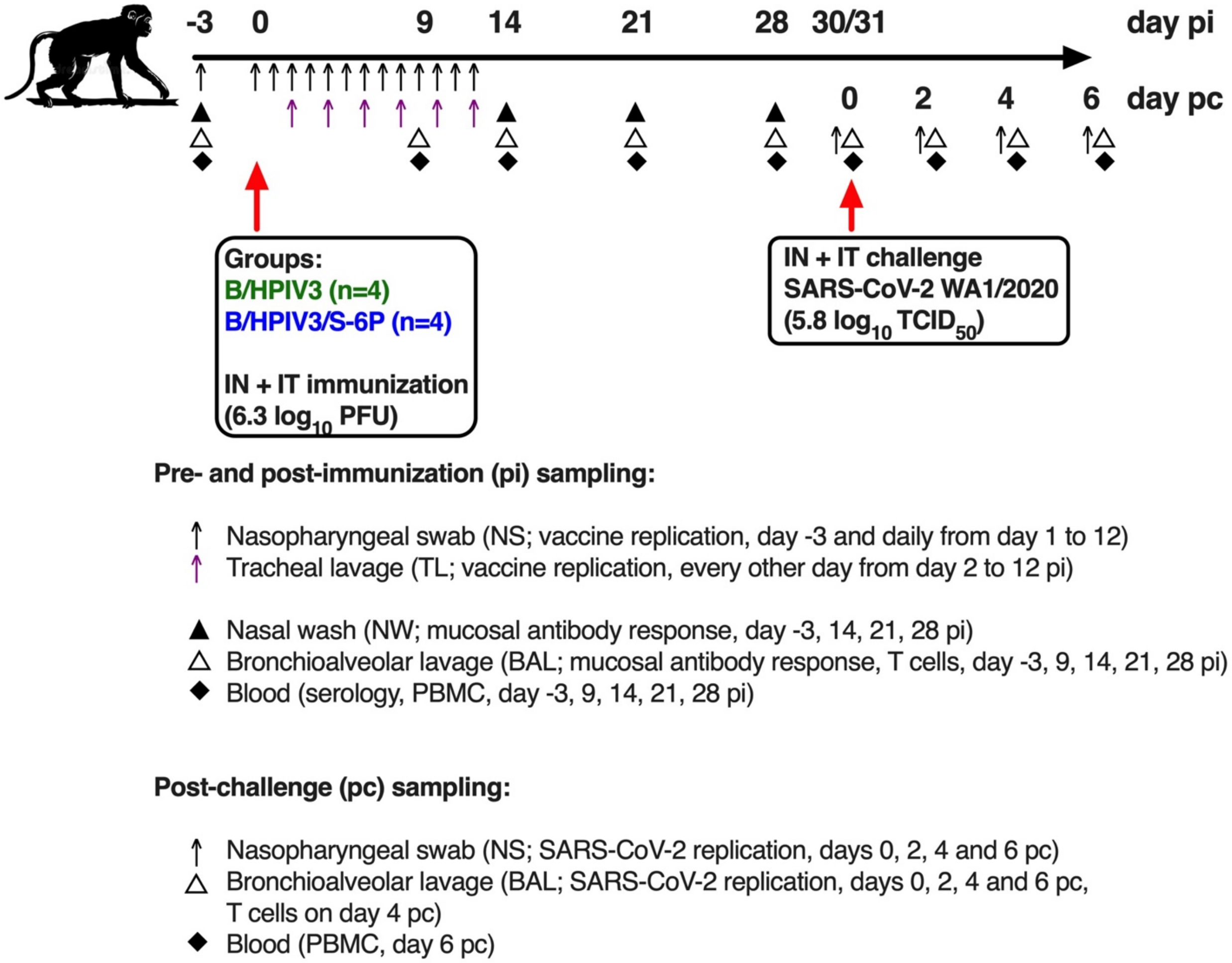


**Figure S1. Timeline of the rhesus macaque study and sampling**
Experimental timeline for the immunization of groups of 4 RMs with the B/HPIV3/S-6P vaccine candidate or the empty B/HPIV3 vector used as a control. Challenge with the SARS-CoV-2 WA1/2020 isolate was performed on day 30 or 31 post-immunization. Pre- and post-challenge sampling schedules are summarized; pi, post-immunization; pc, post challenge. Details are described in materials and methods.

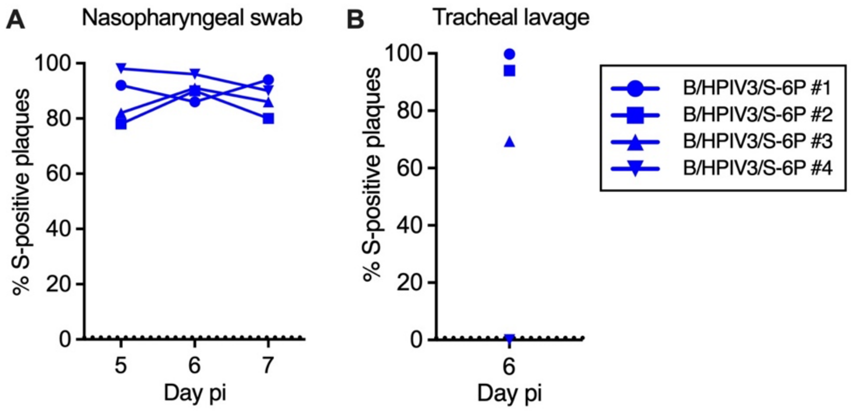


**Figure S2. S expression by B/HPIV3/S-6P in rhesus macaques**
The stability of S expression by B/HPIV3/S-6P in RMs was evaluated by dual-staining immunoplaque assay on Vero cells from nasopharyngeal swab (NS) (**A**) and tracheal lavage (TL) (**B**) samples collected at the peak of vaccine shedding (days 5 through 7). Plaques were immunostained with an HPIV3-specific rabbit hyperimmune serum to detect B/HPIV3 antigens, and a goat hyperimmune serum to the secreted SARS-CoV-2 S to detect co-expression of the S protein, followed by infrared-dye secondary antibodies. Fluorescent staining for PIV3 proteins and SARS-CoV-2 S was visualized in green and red, respectively, generating yellow plaques when merged. The percentage of yellow plaques expressing both HPIV3 and S proteins was determined.

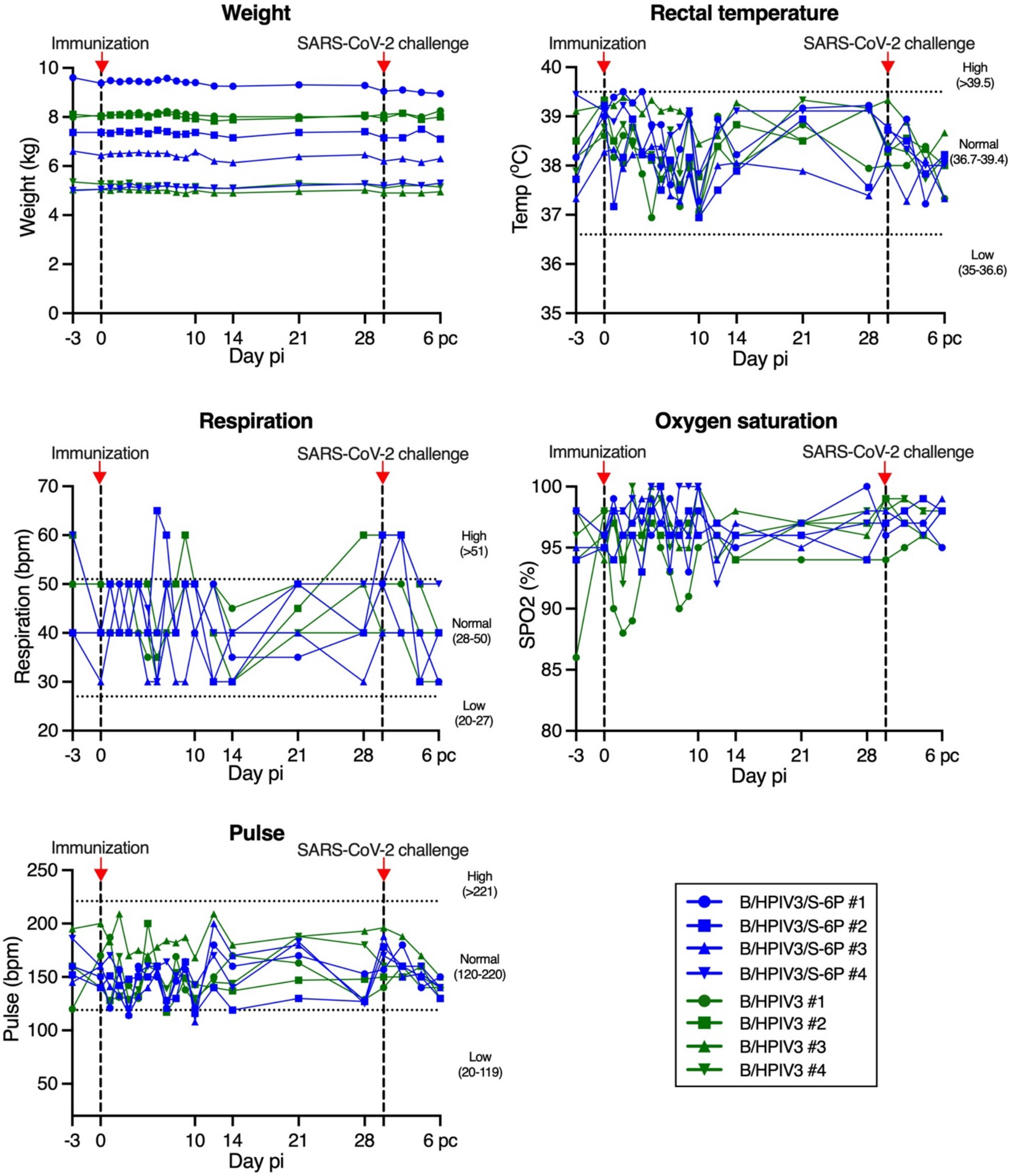


**Figure S3. Vital signs of rhesus macaques after immunization with B/HPIV3 or B/HPIV3/S-6P and SARS-CoV-2 challenge**

Macaques in groups of 4 were immunized with B/HPIV3/S-6P or with the B/HPIV3 (empty vector control). On day 30 post-immunization (pi), animals were challenged in a BSL3 facility with the SARS-CoV-2 WA1/2020 isolate. Animals were euthanized on day 36 pi (day 6 post-challenge). The body weight, rectal temperature, respiration rate, heart rate, and oxygen saturation rate were monitored on the indicated day pi. Timing of immunization and SARS-CoV-2 challenge is indicated by dashed lines and red arrows. pi, post-immunization; pc, post challenge. B/HPIV3/S-6P-immunized animals are represented in blue, and B/HPIV3-immunized animals are represented in green. Each animal is represented by a unique symbol.

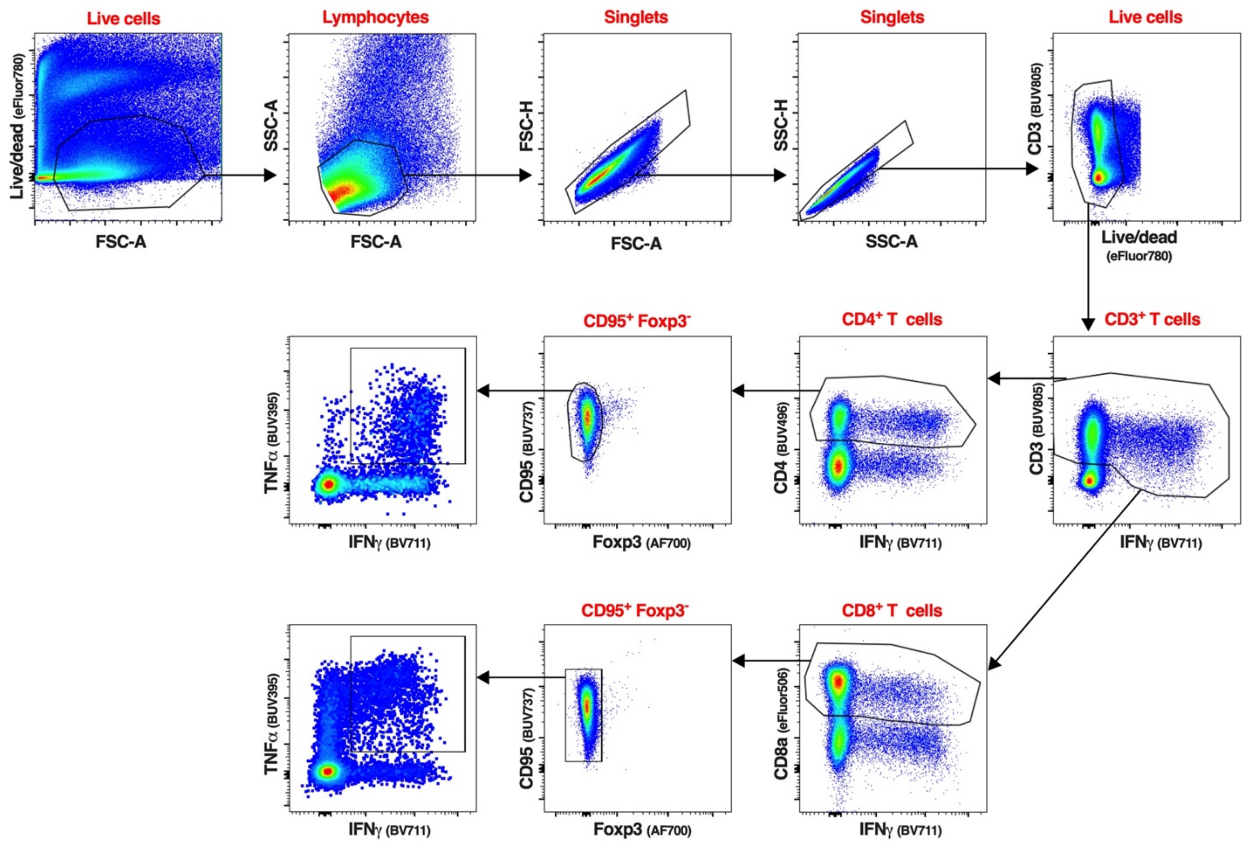


**Figure S4. Gating strategy of the CD4$^+$ and CD8$^+$ T-cells isolated from BAL of rhesus macaques**

Representative flow cytometry dot plots of cells isolated from a BAL sample, visualizing the typical gating strategy used to identify the CD4$^+$ and CD8$^+$ T cell populations described in Figures 4 and 5. The same gating strategy was applied to identify and analyze the CD4$^+$ and CD8$^+$ T-cells from PBMC isolated from the blood (Figures 4 and S5). Live cells were first gated based on a live/dead staining and forward scatter area. Live lymphocytes were identified based on forward and side scatter areas. Then, singlets were selected using a first gate based on forward scatter height and forward scatter area followed by a second gate based on side scatter height and side scatter area. An additional live/dead gating was performed to discard any remaining dead cells. The live single CD3$^+$ IFNγ$^+$ T-cells were next gated using CD3 and IFNγ. As CD3 expression can be downregulated on activated T-cells, a wide CD3 gate was applied. IFNγ$^+$ CD4$^+$ or CD8$^+$ T-cells were next identified using a CD4 or CD8 antibody. Non-naïve, non-regulatory CD4$^+$ or CD8$^+$ T-cells were finally gated using CD95 and Foxp3, respectively. The phenotypic analyses described in Figures 4, 5, S5 and S6 were performed on live single CD3$^+$ CD4$^+$ CD95$^+$ Foxp3$^-$ or live single CD3$^+$ CD8$^+$ CD95$^+$ Foxp3$^-$ T-cells.

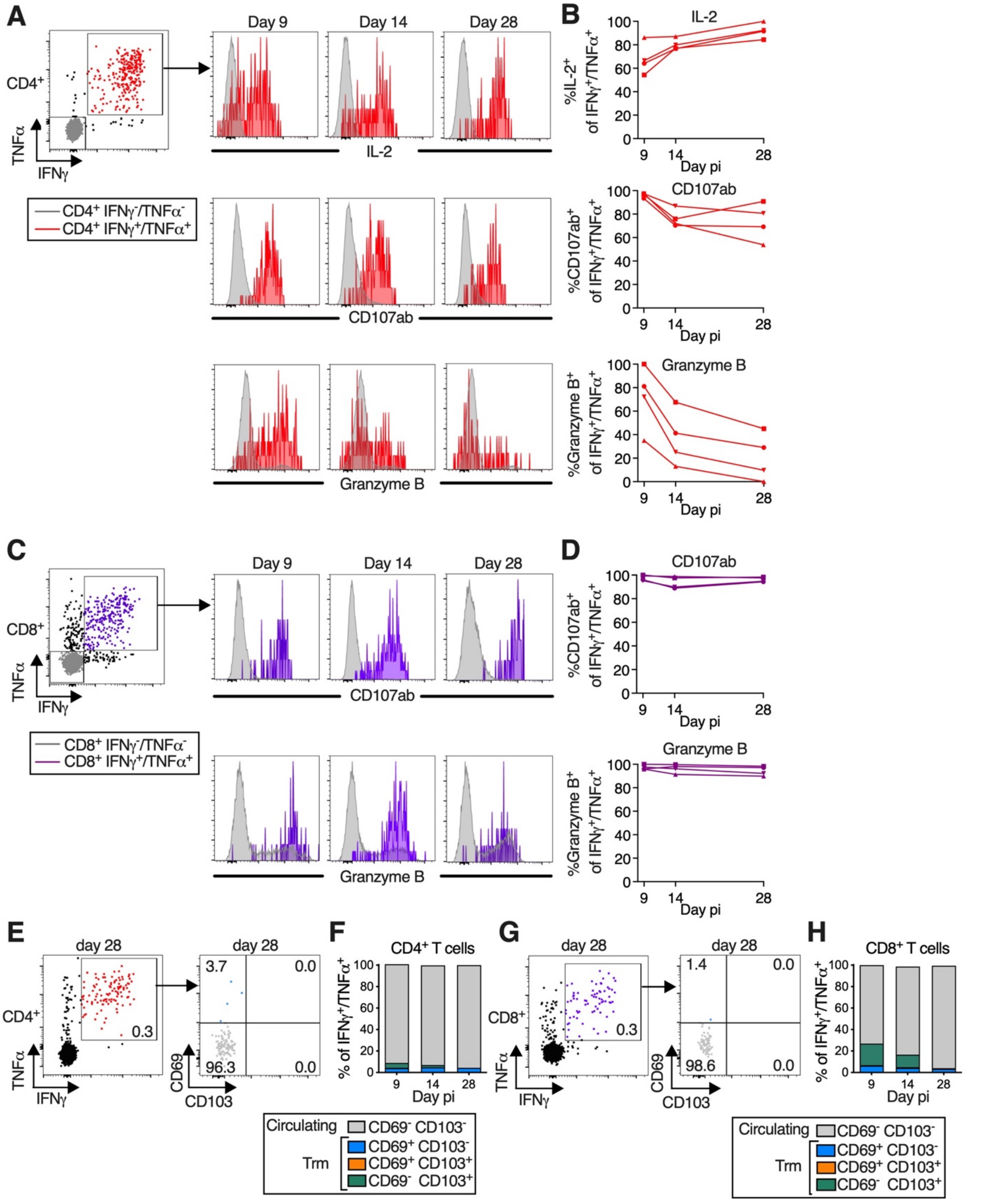


**Figure S5. Phenotype of SARS-CoV-2 S-specific CD4+ and CD8+ T-cells in the blood of the B/HPIV3/S-6P immunized rhesus macaques**

(**A**) Dot blot of the CD4+ T-cells from blood of a representative B/HPIV3/S-6P-immunized RM describing the gating of S-specific IFNγ+/TNFα+ cells (red). The levels of expression of IL-2,

CD107ab and granzyme B by the IFN$\gamma^+$/TNF$\alpha^+$ CD4$^+$ T-cells from the same RM are shown as histograms on the indicated day pi with the IFN$\gamma^+$/TNF$\alpha^+$ CD4$^+$ T-cells (grey) used for reference. (**B**) % of I

FN$\gamma^+$/TNF$\alpha^+$ CD4$^+$ T-cells in the blood of the 4 B/HPIV3/S-6P-immunized RMs that expressed IL-2, CD107ab or granzyme B on the indicated day pi. (**C**) Dot blot of the CD8$^+$ T-cells in the blood of a representative B/HPIV3/S-6P-immunized RM describing the gating of the S-specific IFN$\gamma^+$/TNF$\alpha^+$ cells (purple). The level of expression of CD107ab and granzyme B by the IFN$\gamma^+$/TNF$\alpha^+$ CD4$^+$ T-cells from the same RM are shown as histograms on the indicated day pi with the IFN$\gamma^-$/TNF$\alpha^-$ CD4$^+$ T-cells (grey) used for reference. (**D**) % CD107ab$^+$ or granzyme B$^+$ of IFN$\gamma^+$/TNF$\alpha^+$ CD8$^+$ T-cells on the indicated days pi in the blood of the 4 B/HPIV3/S-6P-immunized RMs. Each macaque is represented by a different symbol. (**E** and **G**) Representative dot plots showing gating on S-specific IFN$\gamma^+$/TNF$\alpha^+$ T-cells (left panels). CD69 and CD103 were used to differentiate circulating (CD69$^-$ CD103$^-$, grey) and tissue-resident memory [Trm; CD69$^+$ CD103$^-$ (blue), CD69$^+$ CD103$^+$ (orange) and CD69$^-$ CD103$^+$ (green)] S-specific IFN$\gamma^+$/TNF$\alpha^+$ T-cells isolated from blood (right panels, % indicated). (**F** and **H**) The median % of the circulating and each of the 3 Trm S-specific IFN$\gamma^+$/TNF$\alpha^+$ CD4$^+$ (**F**) or CD8$^+$ (**H**) T-cell subsets present in blood of 4 B/HPIV3/S-6P-immunized RMs on indicated days are stacked.

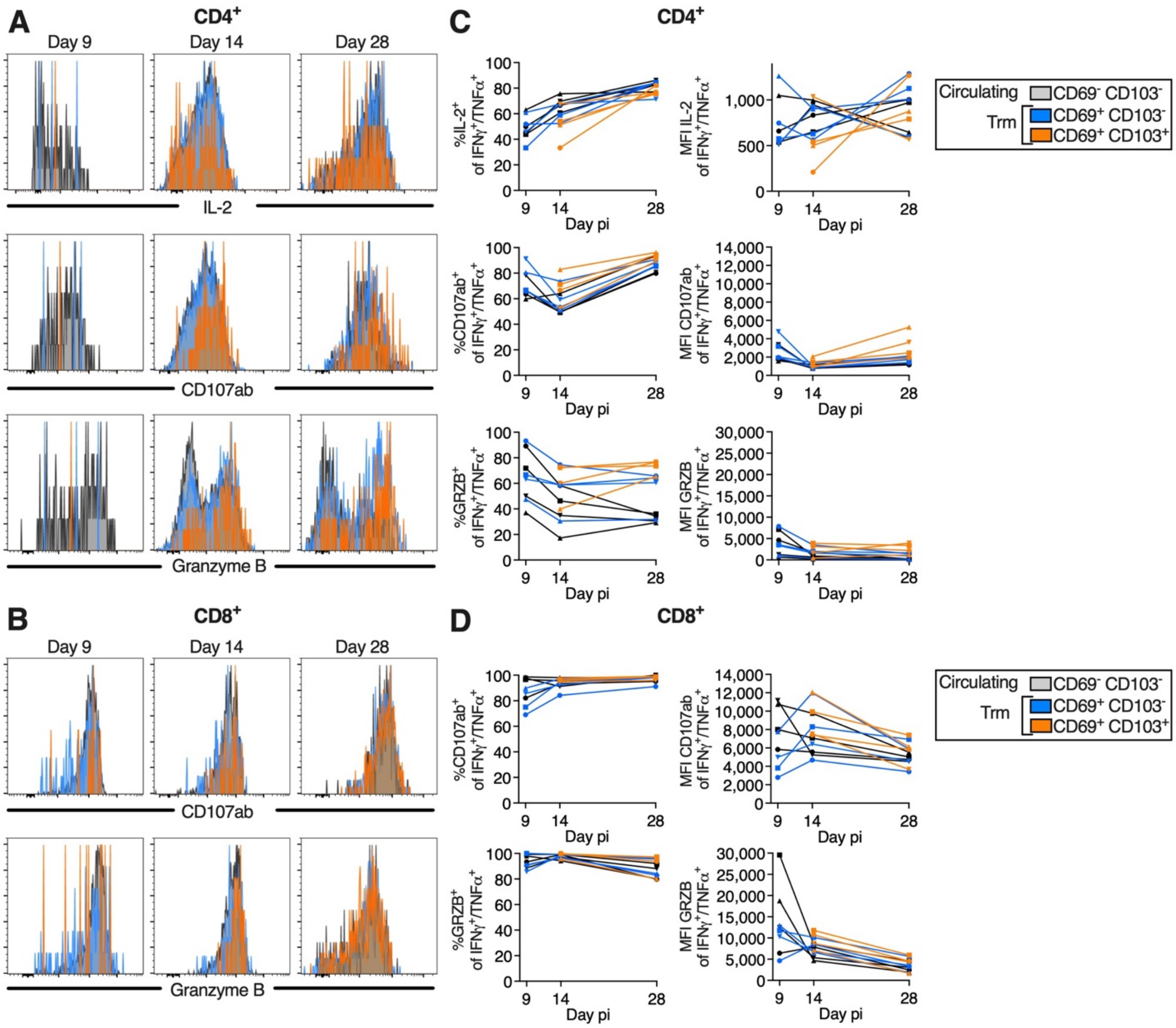


**Figure S6. Comparable phenotype of circulating (CD69$^-$ CD103$^-$) and tissue-resident memory (CD69$^+$ CD103$^-$ and CD69$^+$ CD103$^+$) S-specific IFNγ$^+$/TNFα$^+$ CD4$^+$ and CD8$^+$ T-cells in the airways**

(**A** and **B**) Histograms representing IL2 expression (A only), CD107ab, and granzyme B expression by S-specific circulating and tissue-resident memory (Trm) IFNγ$^+$/TNFα$^+$ CD4$^+$ (A) or CD8$^+$ (B) T-cells obtained from BAL on indicated days pi. (**C** and **D**) % and level of expression (median fluorescence intensity, MFI) of IL2 (CD4$^+$ T-cells only), CD107ab and granzyme B by the S-specific circulating and Trm IFNγ$^+$/TNFα$^+$ CD4$^+$ (C) and CD8$^+$ T-cells (D) in the 4 B/HPIV3/S-6P-immunized RMs. Due to the low frequency of CD69$^+$ CD103$^+$ T-cells on day 9 pi, the frequencies and MFIs of IL-2, CD107ab and granzyme B by this subset are only indicated on days 14 and 28 pi. In C and D, each RM is indicated by a symbol.

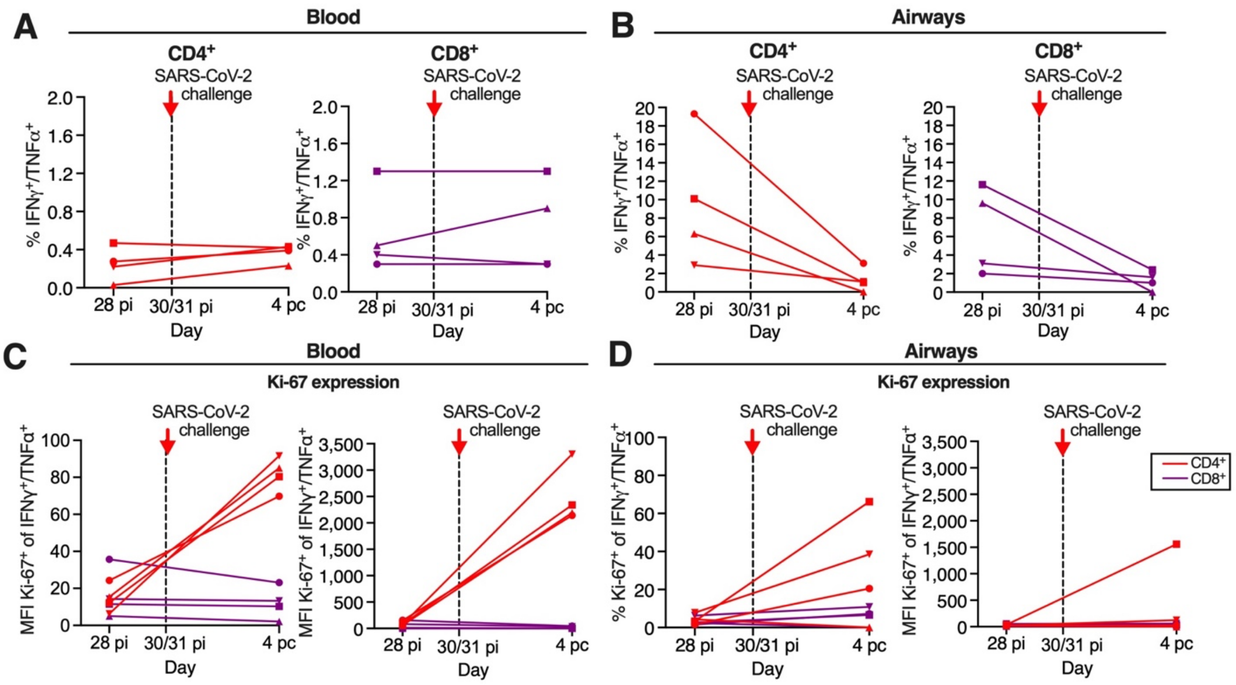


**Figure S7. Expression of proliferation marker Ki-67 by S-specific IFNγ$^+$/TNFα$^+$ CD4$^+$ and CD8$^+$ T-cells 4 days after SARS-CoV-2 challenge**

(**A** and **B**) Background-corrected frequencies of S-specific IFNγ$^+$/TNFα$^+$ CD4$^+$ or CD8$^+$ T-cells from blood (A) or airways (B) on day 28 and 34 pi (equivalent to day 4 post challenge as challenge was performed on day 30 pi). These frequencies are similar to the frequencies shown in Figure 4C, D and Figure 4E, F for the blood and airways, respectively. (**C** and **D**) % and median fluorescence intensity (MFI) of proliferation marker Ki-67 by IFNγ$^+$/TNFα$^+$ CD4$^+$ (red) or CD8$^+$ (purple) T-cells from blood (C) or airways (D) of the 4 B/HPIV3/S-6P-immunized RM, each represented by a different symbol; pi, post-immunization; pc, post challenge

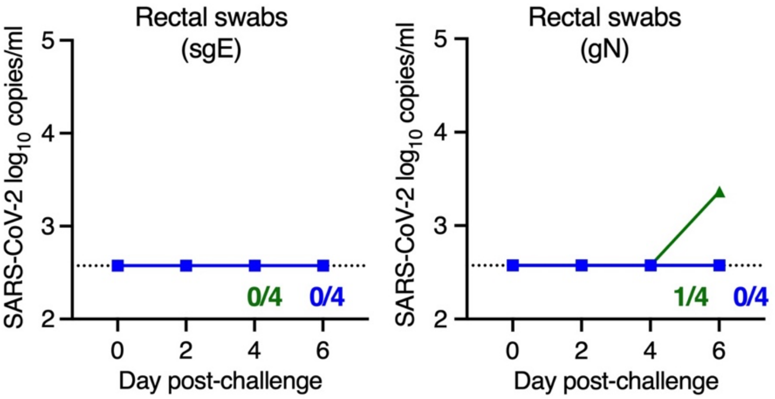


**Figure S8. Quantification of SARS-CoV-2 from rectal swabs**

SARS-CoV-2 subgenomic E (sgE) and genomic N RNA (gN) were quantified by RT-qPCR using RNA extracted from rectal swabs at the indicated day post-challenge. The number of B/HPIV3/S-6P-immunized- or B/HPIV3-immunized RMs with detectable sgE or gN RNA in each set of samples is indicated. The limit of detection was 2.6 $\log_{10}$ copies per ml of rectal swab fluid. B/HPIV3/S-6P-immunized RMs are in blue while B/HPIV3-immunized RMs are in green with each RM indicated by a symbol.
